# Visual Perception of 3D Space and Shape in Time - Part IV - 3D Shape Recognition by 3D Rotation

**DOI:** 10.1101/2022.03.01.482164

**Authors:** Caominh Le, Samantha Pedersen, Nathaniel Chen, Jonathan Chan, Brian Ta, Patrick Wilson, Trevor McCarthy, Emma Barseghyan, Anushka Chauhan, Hind Saif, Jonathan Tu, Darren Wijaya, Annika Zhang, Erica Li, Camille Marangi, Setayesh Nekarae, Felicia Wang, Alice Yanovsky, Umaima Afifa, Javier Carmona, Diego Espino, Leonard Schummer, Phillip Gudijanto, Gurleen Kaur, Andrew Lam, Matthew Mar, Elizabeth Mills, Alexandra Nevins, Elijah Ortiz, Kyle Wheeler, Aaron Blaisdell, Katsushi Arisaka

## Abstract

Human vision has a remarkable ability to recognize complex 3D objects such as faces that appear at any size and 3D orientations at any 3D location. If we initially memorize a face only with a normalized size upfront at the object center, the direct comparison between the one-sized memory and an incoming new image would demand tremendous mental frame translations in 7D. How can we perform such a demanding task so promptly as we experience it in our daily lives?

This paper specifically addresses the recognition of human faces with arbitrary 3D orientation in the [Roll, Yaw, Pitch] axes. According to our new model of **NHT** (Neural Holography Tomography), space is represented by time utilizing the phase of the alpha brainwave. This principle should be applicable to any mental rotation in 3D; thus, it predicts that extra time is required to perceive a rotated face to revolve it back to upright by the constant-speed alpha wave.

To assess this hypothesis, we designed a reaction time (RT) experiment, where participants were first asked to memorize sets of upright unfamiliar faces. Following the memorization phase, similar stimuli with a wide range of rotating faces in 3D were presented, and RTs were recorded. As expected, the memorized upfront face was the fastest RT. The excess of the RT was observed proportional to the rotating angle in all [Roll, Yaw, Pitch] axes. Roll had the flattest slope, whereas upper Pitch was the steepest. We suspect that Roll is the swiftest mental operation because it can be conducted by the linear frame translation on the log-polar retinotopy of the visual cortex.

## 1 Introduction

Most humans utilize visual input as their primary source of sensory information to understand and respond to their surroundings. Past research has mapped the complex network of brain structures that convey and process this information to at least thirty different cortical areas (Pessoa, 2014). Facial recognition, the ability to identify the face of someone familiar, is particularly key to facilitating everyday social interactions (S. Favelle & Palmisano, 2018). In fact, it is so fundamental that human infants are able to instantly discriminate between familiar and unfamiliar faces at various angles (Turati et al., 2008).

Under the two-streams hypothesis, visual processing can be defined by two distinct pathways in the brain; dorsal and ventral (Goodale & Milner, 1992). Although recent evidence has shown that visual processing may not be as binary (Zachariou et al., 2014), within the two-streams hypothesis, the dorsal pathway constitutes the ‘where’ pathway of spatial perception and processes locational and orientational information, while the ventral pathway constitutes the ‘what’ pathway of object perception and facial recognition. Moreover, the dorsal pathway has been shown to process information and guide actions without accompanying conscious knowledge, while the ventral pathway is involved in conscious perception (Fang & He, 2005). In the commonly cited ventral pathway model, visual information proceeds from the primary visual cortex (V1) through the anterior inferotemporal (aIT) cortex, where the average receptive field size and latency increases (Kravitz et al., 2013). V1 orientation vectors are only weakly selective due to receiving input solely from the lateral geniculate nucleus (LGN) (McLaughlin et al., 2000), but become more strongly orientationally selective throughout the pathway such as in V3, where a large majority of neurons are orientationally selective (Felleman & Van Essen, 1987).

It has been long supported that semantic information is processed in the ventral pathway by a log-polar coordinate system in humans and primates (Arcaro et al., 2009; Engel et al., 1997; Kolster et al., 2014). However, the reason for this distortion on the visual cortex has not been fully understood in the context of object recognition, especially with the failure of the current bottom-up theory of visual processing to address the efficiency and accuracy of human facial recognition. A top-down signal connecting visual input and internal imagery is necessary to allow rapid recognition of familiar faces at any size, location, or rotation in as little as 200 ms (Caharel et al., 2015; Dijkstra et al., 2017). Formed from contextual reinforcement, top-down pattern recognition also allows for efficient analysis and categorization of key facial features (Puce et al., 1999).

The viewpoint dependence of facial recognition on Roll, Yaw, and Pitch has been previously studied, and performance has been found to be most optimal about the Roll axis, followed by Yaw and Pitch (S. K. Favelle et al., 2011). Another study found that participants assigned to a front or ¾ yaw comparison view performed better than those assigned to pitch-up or pitch-down views, probably due to how these angles maximize the contour and projection of feature information, allowing better extraction of 3D information (S. Favelle & Palmisano, 2018). In general, faces are processed holistically, but when skewed at different angles, the visual processing mechanism of the cortex is forced to focus on individual key features to recognize the face (S. Favelle & Palmisano, 2012). Ultimately, holistic processing requires significantly more time than digesting a few key features but may lead to better accuracy.

We have recently proposed the new concepts of MePMoS (Memory-Prediction-Motion-Sensing) and NHT (Neural Holographic Tomography) to map the processes of this conscious top-down approach (Arisaka, 2022a, 2022b; Arisaka & Blaisdell, 2022). Facial recognition occurs when a relevant memorized pattern of the stimulus undergoes 7 degrees of transformation to overlap with the incoming image, and this transformation can be expressed as measurements in the latency of alpha brainwaves. In the past, alpha oscillations have been studied as a possible mechanism to encode visual input by inhibiting irrelevant brain structures and directing information to appropriate neural structures for semantic encoding during early perception or semantic retrieval during object recognition (Klimesch et al., 2011). Under the MePMoS model, we predict that alpha waves are also involved in scaling and rotating memorized data through variable phase shifts determined by a time delay to overlap incoming signals and allow recognition. After the initial visual input is received an alpha wave cycle, the relevant memory is extracted and transformed by alpha wave phase shifts to predict the appearance of the incoming stimulus. Upon overlap, two more alpha cycles are required for conscious awareness of the congruency and instructions for motion output, placing the simplest conscious recognition and behavior at around 400 ms.

The NHT model also better explains the significance of the log-polar distortion of the ventral pathway. Conversion of incoming Cartesian images to log-polar makes the projection onto the V1 visual cortex scale and roll-rotation invariant. Rescaling the memorized image, causing key features to move to or from the periphery, becomes a simple horizontal translation of the distorted projection on the log axis while rotating the stimulus along the Roll axis becomes a simple vertical translation on the polar coordinate axis. This translation can be encoded through phase shifts of top-down signals on the timescale of constant-frequency alpha waves. Larger rotations about the roll axis require larger translations in the projected V1 image, leading to higher phase shifts in alpha waves and time delays on the order of 100 ms.

Unlike Roll, Yaw and Pitch involve more extensive stepwise transformations to overlap memorized data to incoming stimuli. The brain must first translate the memory must first from an egocentric frame of reference to an object-centric frame. Once centered at the object, the memory can then be rotated along the yaw or pitch axes. Finally, the frame can be restored to the original egocentric frame of reference and compared to incoming stimuli from the same frame. While all three steps may require shifts in brain wave patterns, only the rotation step differs between stimuli, given that they were memorized at the same distance and location. This transformation can be observed by reaction time differences, with larger rotations leading to higher alpha wave phase shifts and latencies about 100 ms apart.

## 2 Results

Data from the four independent groups was analyzed in a combined data pool and separately. While each group had differences in their protocols, testing environment, and knowledge of the research, there was an impressive agreement between corresponding protocols that justified analyzing all groups in a combined data pool. Within each protocol, data was divided into negative and positive rotation angles. Negative angles consisted of counterclockwise Roll, leftward Yaw, and downward Pitch rotations. Positive angles consisted of clockwise Roll, rightward Yaw, and upward Pitch rotations. 0 and 180 degrees (for Roll) were analyzed in both data halves.

From each individual participant’s data, average reaction times and standard errors were calculated for each angle of rotation, and linear fits were calculated for positive and negative angles using minimum chi-square estimation. A histogram of reduced chi-square values from both positive and negative trendlines was generated alongside a gamma distribution. Gamma distribution modes near 1.00 suggested that most individual datasets could be modeled linearly and provided evidence that the aggregate pool could also be modeled linearly. Average reaction times were calculated from the aggregated data pool. Standard errors were calculated after normalizing the data around the aggregate mean by Cousineau normalization (Morey, 2008). Using these parameters, linear fits were calculated for positive and negative angles using minimum chi-square estimation, and the reduced chi-square was used to determine the strength of the fit. Reduced chi-squares greater than one suggested the possibility of more optimal non-linear fits while models with reduced chi-square less than one were judged based on the size of error bars. Plots showing stacked individual participant data before and after normalization can be found in the Supplementary Figures along with reduced chi-square histograms. An example of data analysis plots is shown in Figure 2.

**Figure 1.**
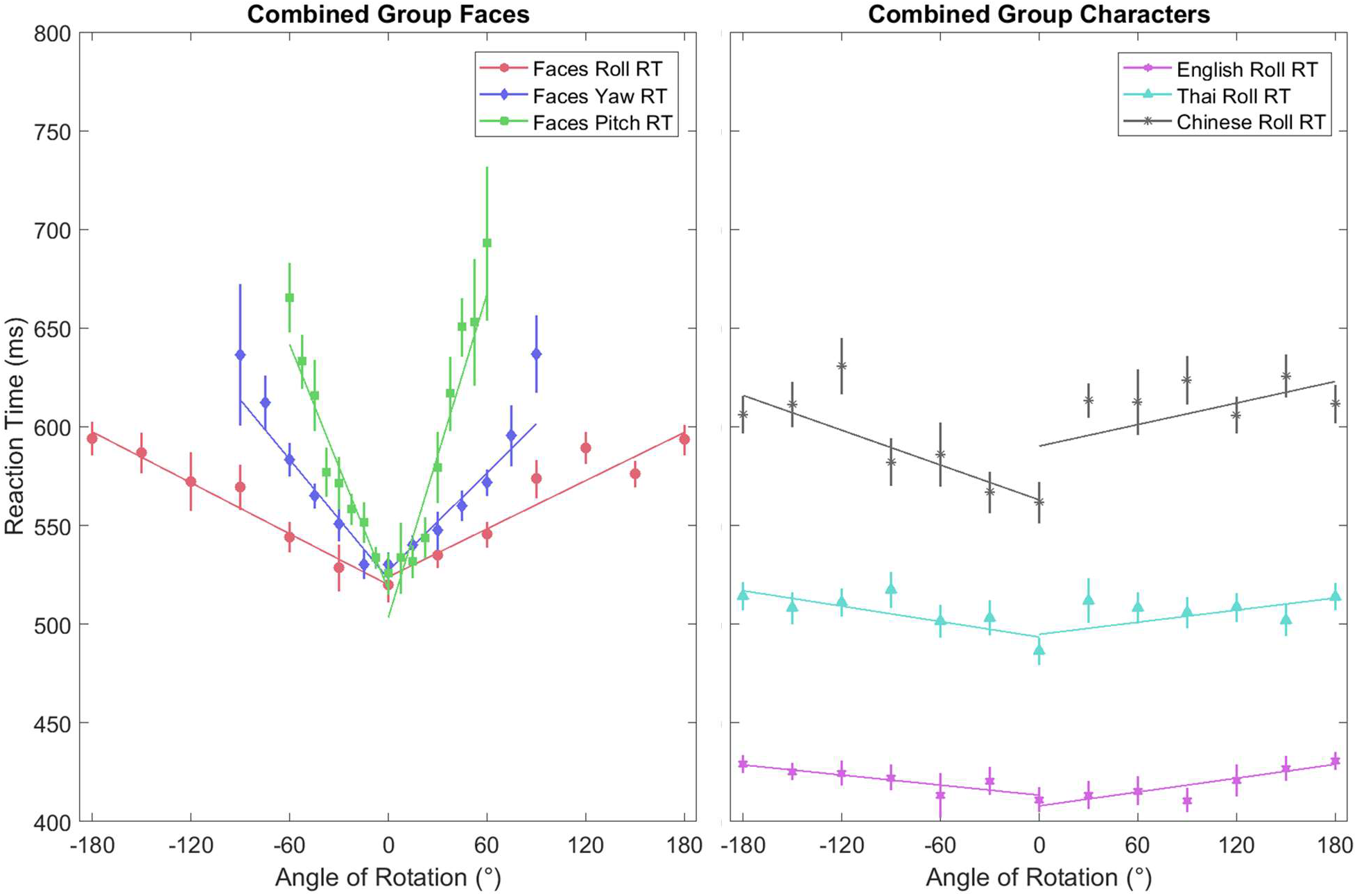
Combined Group data showing face and character protocols on a common axis. Among the faces, Roll was the most symmetric and least steep while Pitch was the most asymmetric and the steepest. Among characters, English and Thai were almost flat with Thai having a longer central reaction time while Chinese had the longest reaction times and was almost as steep as Face Roll.

**Figure 2.**
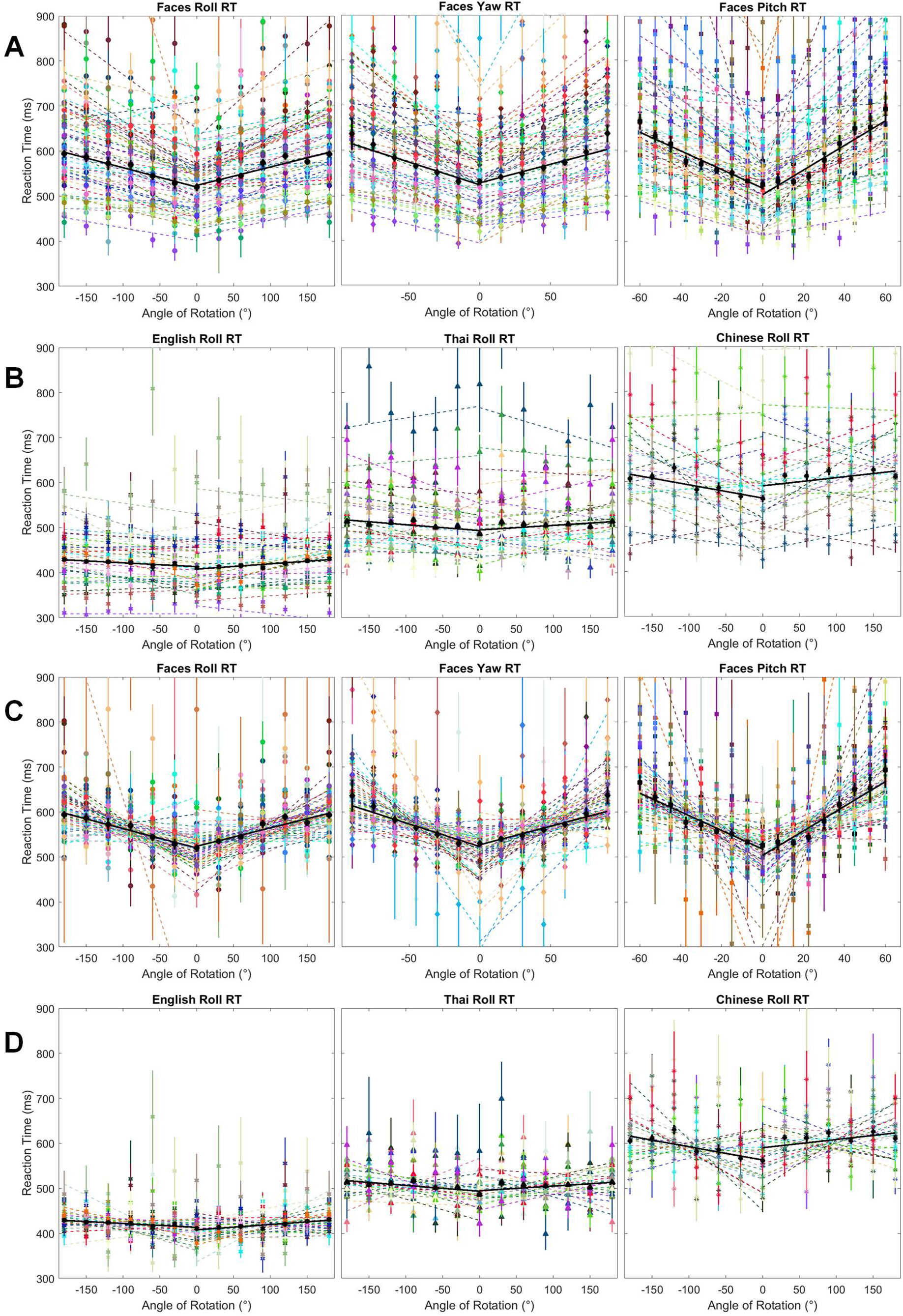
Supplementary data analysis plots for combined data pool. (A) Raw reaction times (RT) for faces against roll, yaw, and pitch rotations. Error bars were calculated after a Cousineau normalization was applied. (B) Raw RTs for English, Thai, and Chinese against roll rotation. Error bars were calculated after a Cousineau normalization was applied. (C) Normalized RTs for face protocols. (D) Normalized RTs for character protocols.

### 2.1 Combined Analysis

All groups were analyzed together as a single aggregate group (Figure 1). Differences and consistencies between groups are considered in subsequent sections.

**Table 1.** Significant parameters for combined analysis. Parameters for separate groups are shown in **Table 3** and Supplementary Table S1 and S2. Individual reduced chi-square modes were calculated from gamma distribution curves of individual participant reduced chi-squares for right and left trendlines. (*) The distribution of reduced chi-squares for Thai Roll appeared uniformly distributed between 0 and 3 and lacked a clear peak although one was calculated from a gamma fit.

| Experiment Information |  |  | Negative (Counterclockwise, Left, Down) |  |  |  | Positive (Clockwise, Right, Up) |  |  |  |
| --- | --- | --- | --- | --- | --- | --- | --- | --- | --- | --- |
| Protocol | n | Individual Reduced $\chi^2$ Modes | Slope (ms/°) | Intercept (ms) | R | Reduced $\chi^2$ | Slope (ms/°) | Intercept (ms) | R | Reduced $\chi^2$ |
| Face Roll | 56 | 1.10 | -0.43 | 521 | -0.99 | 0.25 | 0.40 | 525 | 0.94 | 1.68 |
| Face Yaw | 56 | 0.92 | -1.01 | 522 | -0.98 | 0.81 | 0.84 | 526 | 0.95 | 1.02 |
| Face Pitch | 54 | 1.39 | -2.07 | 517 | -0.97 | 0.85 | 2.73 | 503 | 0.97 | 2.14 |
| English Roll | 30 | 0.99 | -0.09 | 426 | -0.88 | 0.26 | 0.12 | 420 | 0.89 | 0.63 |
| Thai Roll | 25 | 1.03* | -0.13 | 498 | -0.77 | 1.13 | 0.10 | 500 | 0.53 | 1.47 |
| Chinese Roll | 22 | 1.19 | -0.29 | 563 | -0.82 | 2.07 | 0.18 | 590 | 0.60 | 4.77 |

#### Unfamiliar Faces Rotation

All three face protocols resulted in similar average y-intercepts and V-shapes with Roll being the least steep and Pitch being the steepest. Roll had a symmetric V-shape while Yaw had a slightly asymmetric V-shape and Pitch had a very asymmetric V-shape. All three protocols had small reduced-chi-square values for negative angles, indicating strong linear fits. Interestingly, the reduced chi-square values for positive angles were greater, indicating weaker but acceptable linear fits. Both roll and yaw deviated from the linear trendline at the higher angles with a slight decrease in reaction time around ±150° roll and an upward curve towards ±90° yaw. In the pitch protocol, there was a downward deviation from the linear prediction at 22.5°.

#### Character Roll

Both English and Thai roll protocols were symmetric and almost flat, being four times less steep than faces roll. Thai was parallel to English and took on average 85 - 90 ms longer at any given angle. Compared to faces roll, participants responded 25 seconds faster to unrotated Thai than unrotated faces. Both linear predictions were strong fits as indicated by small reduced-chi-squares, especially for negative angles. For Chinese Roll, there was a weaker fit with a large reduced-chi-square, suggesting that the relationship between reaction time and rotation was non-linear. Additionally, the central reaction time was 40 - 60 ms slower than for unrotated faces. The trendlines were noncontinuous with a large jump in reaction time from 0° to +30°. The data for negative angles had a slope between that of character roll and face roll while the data for positive angles was flatter and more closely resembled character data. As with faces roll, there was a drop-in reaction time around ±150° for both Thai and Chinese.

**Table 2.** Gamma Distribution modes for Reduced Chi-squares of linear fits to individual participant data. (*) Some distributions were more uniformly distributed, skewed, or had an additional mode. For group A, English Roll was bimodal, and Thai Roll was skewed, giving it a smaller calculated mode of 1.33 than the estimated actual mode. For group B, Thai Roll was bimodal while Chinese Roll had a lower calculated mode of 1.30 than the estimated actual mode.

| Protocol | Group | n | Individual Reduced $\chi^2$<br>Modes | Protocol | Group | n | Individual Reduced $\chi^2$<br>Modes |
| --- | --- | --- | --- | --- | --- | --- | --- |
| Face Roll | A | 14 | 1.04 | English Roll | A | 14 | 1.15, 3* (bimodal) |
|  | B | 15 | 1.18 |  | B | 15 | 1.02 |
|  | C | 13 | 0.94 |  | C | 13 | - |
|  | D | 14 | 1.31 |  | D | 14 | 0.89 |
| Face Yaw | A | 14 | 0.47 | Thai Roll | A | 14 | 2.7* |
|  | B | 15 | 1.04 |  | B | 15 | 0.70, 2.8* |
|  | C | 13 | 1.17 |  | C | 13 | - |
|  | D | 14 | 0.96 |  | D | 14 | 1.33 |
| Face Pitch | A | 12 | 1.66 | Chinese Roll | A | 12 | - |
|  | B | 15 | 1.76 |  | B | 15 | 2* |
|  | C | 13 | 1.23 |  | C | 13 | - |
|  | D | 14 | 0.67 |  | D | 14 | 1.11 |

**Table 3.**
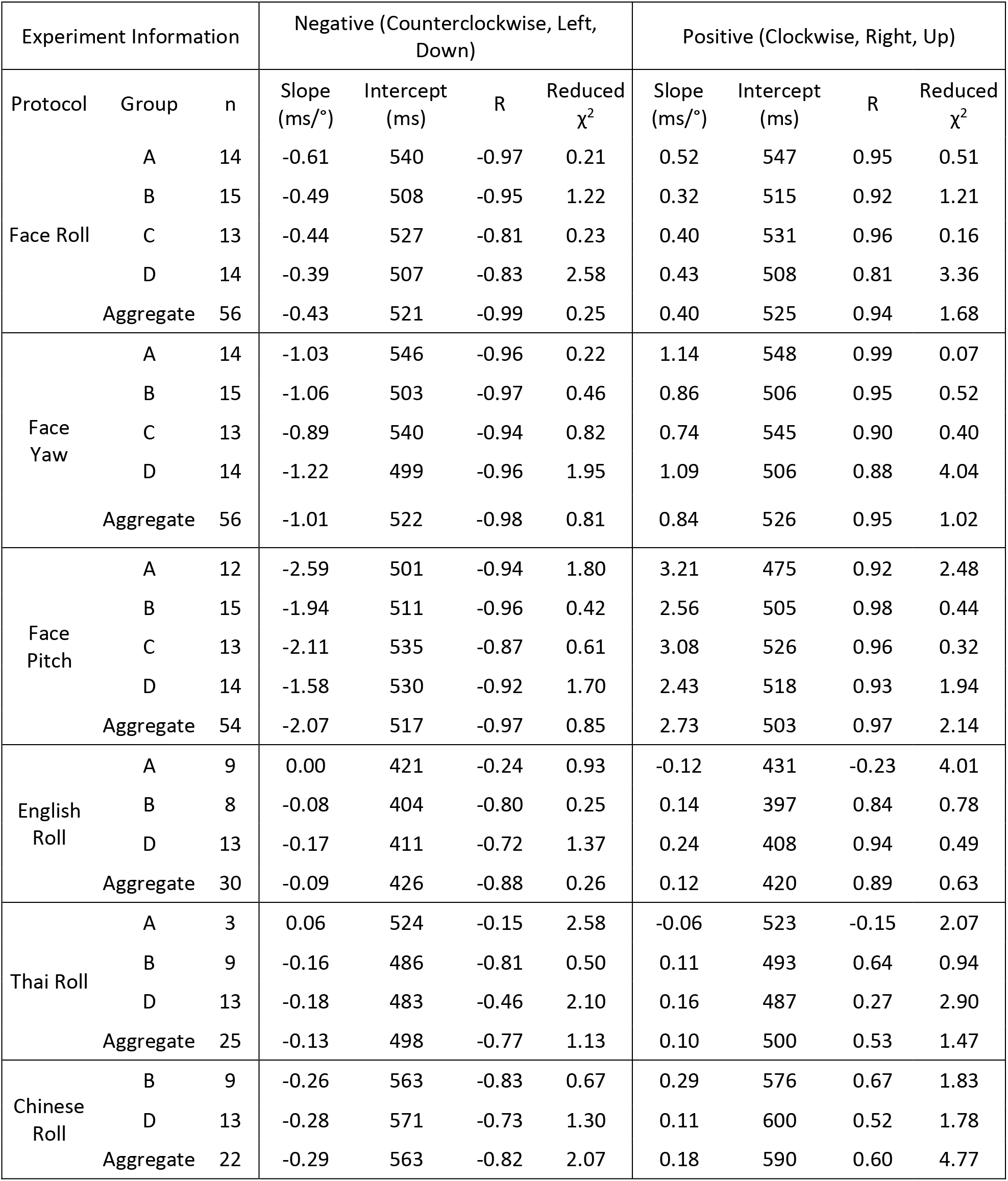
Significant parameters for unfamiliar face rotations and character roll. A complete table of parameters with corresponding error ranges can be found in Supplementary Table S1 and S2.

| Experiment Information |  |  | Negative (Counterclockwise, Left, Down) |  |  |  | Positive (Clockwise, Right, Up) |  |  |  |
| --- | --- | --- | --- | --- | --- | --- | --- | --- | --- | --- |
| Protocol | Group | n | Slope (ms/°) | Intercept (ms) | R | Reduced $\chi^2$ | Slope (ms/°) | Intercept (ms) | R | Reduced $\chi^2$ |
| Face Roll | A | 14 | -0.61 | 540 | -0.97 | 0.21 | 0.52 | 547 | 0.95 | 0.51 |
|  | B | 15 | -0.49 | 508 | -0.95 | 1.22 | 0.32 | 515 | 0.92 | 1.21 |
|  | C | 13 | -0.44 | 527 | -0.81 | 0.23 | 0.40 | 531 | 0.96 | 0.16 |
|  | D | 14 | -0.39 | 507 | -0.83 | 2.58 | 0.43 | 508 | 0.81 | 3.36 |
|  | Aggregate | 56 | -0.43 | 521 | -0.99 | 0.25 | 0.40 | 525 | 0.94 | 1.68 |
| Face Yaw | A | 14 | -1.03 | 546 | -0.96 | 0.22 | 1.14 | 548 | 0.99 | 0.07 |
|  | B | 15 | -1.06 | 503 | -0.97 | 0.46 | 0.86 | 506 | 0.95 | 0.52 |
|  | C | 13 | -0.89 | 540 | -0.94 | 0.82 | 0.74 | 545 | 0.90 | 0.40 |
|  | D | 14 | -1.22 | 499 | -0.96 | 1.95 | 1.09 | 506 | 0.88 | 4.04 |
|  | Aggregate | 56 | -1.01 | 522 | -0.98 | 0.81 | 0.84 | 526 | 0.95 | 1.02 |
| Face Pitch | A | 12 | -2.59 | 501 | -0.94 | 1.80 | 3.21 | 475 | 0.92 | 2.48 |
|  | B | 15 | -1.94 | 511 | -0.96 | 0.42 | 2.56 | 505 | 0.98 | 0.44 |
|  | C | 13 | -2.11 | 535 | -0.87 | 0.61 | 3.08 | 526 | 0.96 | 0.32 |
|  | D | 14 | -1.58 | 530 | -0.92 | 1.70 | 2.43 | 518 | 0.93 | 1.94 |
|  | Aggregate | 54 | -2.07 | 517 | -0.97 | 0.85 | 2.73 | 503 | 0.97 | 2.14 |
| English Roll | A | 9 | 0.00 | 421 | -0.24 | 0.93 | -0.12 | 431 | -0.23 | 4.01 |
|  | B | 8 | -0.08 | 404 | -0.80 | 0.25 | 0.14 | 397 | 0.84 | 0.78 |
|  | D | 13 | -0.17 | 411 | -0.72 | 1.37 | 0.24 | 408 | 0.94 | 0.49 |
|  | Aggregate | 30 | -0.09 | 426 | -0.88 | 0.26 | 0.12 | 420 | 0.89 | 0.63 |
| Thai Roll | A | 3 | 0.06 | 524 | -0.15 | 2.58 | -0.06 | 523 | -0.15 | 2.07 |
|  | B | 9 | -0.16 | 486 | -0.81 | 0.50 | 0.11 | 493 | 0.64 | 0.94 |
|  | D | 13 | -0.18 | 483 | -0.46 | 2.10 | 0.16 | 487 | 0.27 | 2.90 |
|  | Aggregate | 25 | -0.13 | 498 | -0.77 | 1.13 | 0.10 | 500 | 0.53 | 1.47 |
| Chinese Roll | B | 9 | -0.26 | 563 | -0.83 | 0.67 | 0.29 | 576 | 0.67 | 1.83 |
|  | D | 13 | -0.28 | 571 | -0.73 | 1.30 | 0.11 | 600 | 0.52 | 1.78 |
|  | Aggregate | 22 | -0.29 | 563 | -0.82 | 2.07 | 0.18 | 590 | 0.60 | 4.77 |

**Figure 3.**
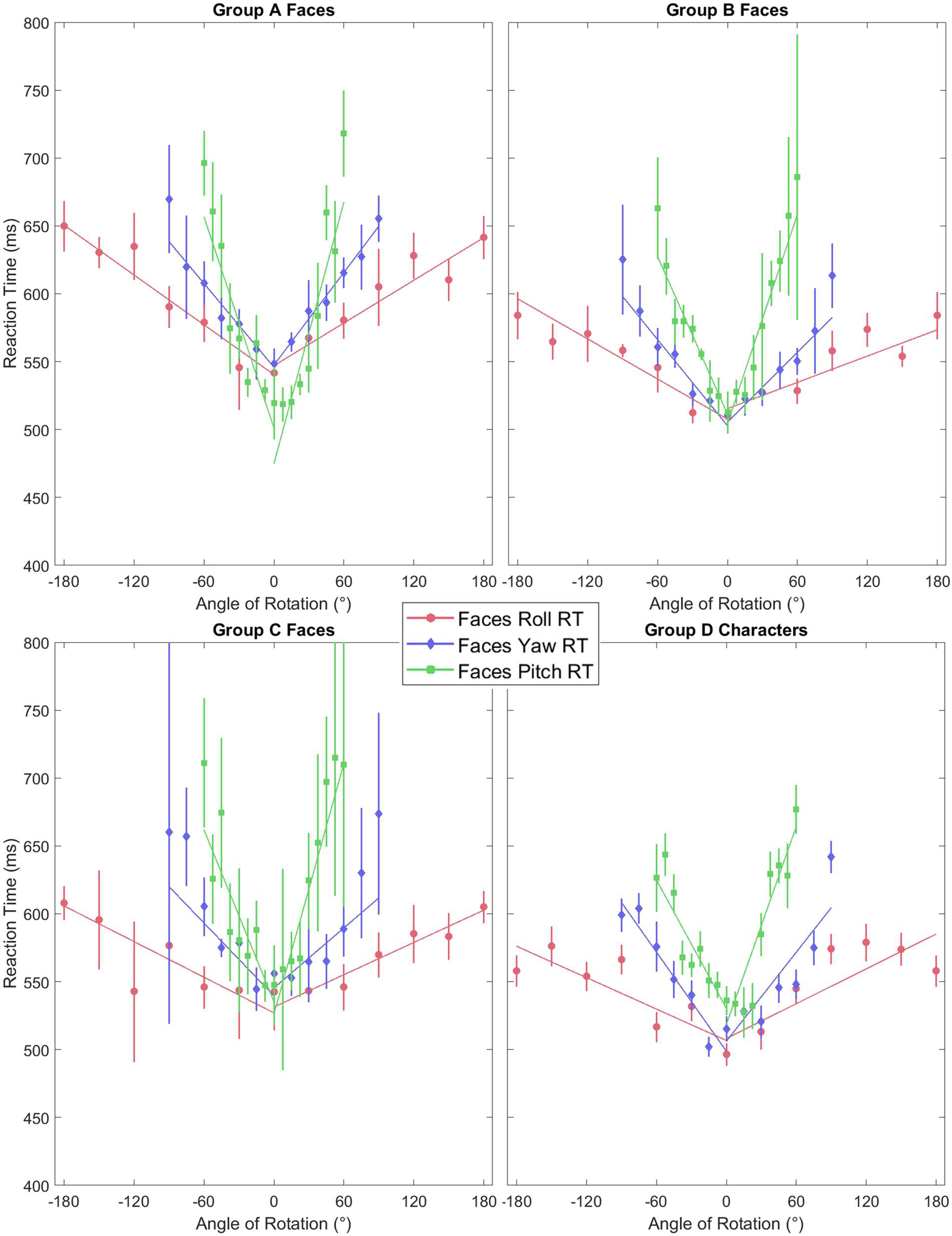
Linear fits to aggregate Faces data for Groups A, B C, and D. Raw and normalized protocol plots along with reduced chi-square histograms are included in the Supplementary Figures.

**Figure 4.**
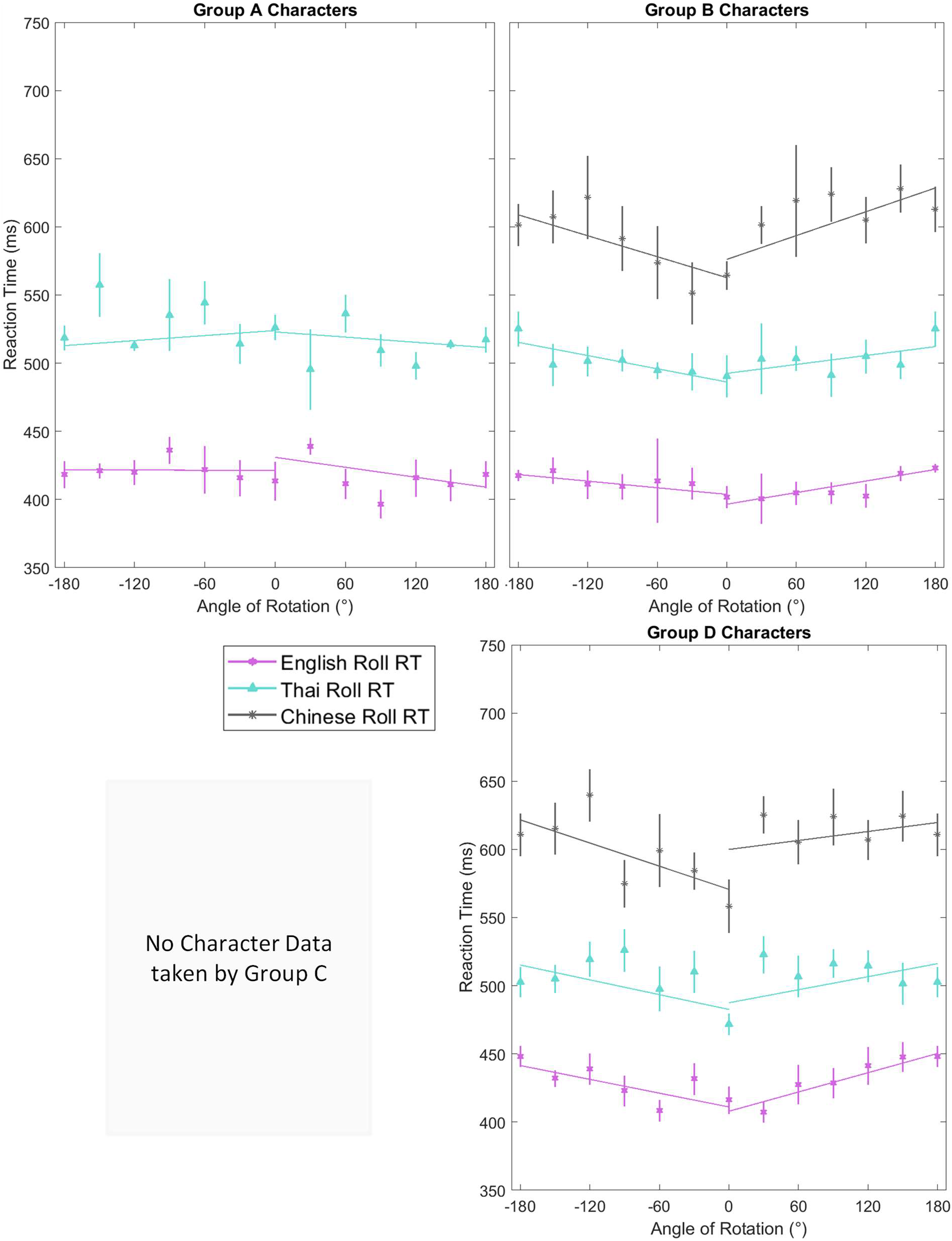
Linear fits to aggregate Characters data for Groups A, B C, and D. Raw and normalized protocol plots along with reduced chi-square histograms are included in the Supplementary Figures.

### 2.2 Separate Group Analysis

#### Internal Group A

Internal Group A took reaction time data through attractor vs. distractor tests. Gamma distribution modes for Roll and Yaw were near or less than 1, providing strong support for a linear relationship between reaction time and rotational angle, while the mode for Pitch was higher but acceptable. Roll (n = 14) was nearly symmetric while Yaw (n = 14) was more symmetric, and Pitch (n = 12) was more asymmetric. As with the aggregate, roll was the least steep and pitch was the steepest protocol. After normalizing individual datasets about the aggregate via Cousineau normalization, roll and yaw appeared to have strong linear fits with small reduced-chi-squares and reasonable error bars. However, pitch had a larger reduced chi-square, suggesting that the relationship may not be completely linear, especially at central angles where the data appears to curve like a parabola. The pitch data was downshifted from the other protocols with a faster unrotated reaction time.

For English (n = 9), the gamma distribution was bimodal with a larger mode near 1 and a smaller mode near 3. Hence, while most participants followed linear reaction time changes, it is possible that some had nonlinear reaction time changes. Thai (n = 3) Roll had a single mode near 2.7, suggesting a nonlinear relationship. Nevertheless, there was little correlation between reaction time and degree of rotation, as linear predictions for both appeared flat and had inverted V-shapes. As with the aggregate, participants tended to take longer to recognize Thai than English. Thai data had more variation between data at each angle than English, leading to a weaker linear prediction and larger reduced chi-square. English had a stronger linear fit with a reduced chi-square near one.

#### Internal Group B

Internal Group B took reaction time data through 3-choice reaction time tests. As with group A, face data gamma distribution modes supported linear predictions for Roll and Yaw, while the mode for Pitch was higher. Both roll (n = 15) and yaw (n = 15) were slightly asymmetric with steeper negative slopes than positive slopes. Pitch (n = 15) was asymmetric in the opposite direction, with a steeper positive slope than negative slope. Nonetheless, groups A and B agreed with roll being the least steep and pitch being the steepest. Roll had the strongest linear prediction with a reduced chi-square near one while yaw and pitch had large error bars at larger rotation angles due to a subset of participants having nonlinear relations between the reaction time and rotation or variation in linear slope between participants, leading to small reduced-chi-squares.

For character roll, English (n = 8) and Chinese (n = 9) both had single gamma distribution modes near 1 and 2 respectively, suggesting that Chinese may not be strongly linear. Thai (n = 9) was bimodal with most participants having linear relationships between reaction time and rotation and a smaller group having nonlinear relationships. In contrast to Group A, the aggregated analysis for English and Thai resulted in shallow but upright V-shapes with stronger correlations and smaller reduced chi-squares. There appeared to be a 20 ms shift in reaction time between the unrotated and fully rotated characters. For Chinese, the slopes appeared intermediate between the simpler characters and the face roll, but there was a notable nonlinear curve to the data that plateaued at higher rotations. Hence, Chinese roll recognition may be modeled more optimally by a non-linear regression.

#### External Group C

External Group C took reaction time data for only faces through shortened 3-choice reaction time tests. Gamma distribution modes supported linear predictions for all three protocols. Roll (n = 13) was symmetric and had the least steep slope. Like group B, yaw (n = 13) was slightly asymmetric with a steeper negative slope than positive slope, and Pitch (n = 13) had the steepest slopes, which were asymmetric with a steeper positive slope than negative slope. Except for data at −120°, roll had reasonable error bars and small reduced chi-squares, indicating a strong linear fit. On the other hand, yaw and pitch had very large error bars due to variation in slopes between participants, leading to decreased reduced chi-squares. Pitch appeared linear, but yaw appeared to curve upward from the linear prediction at larger rotations.

#### External Group D

External Group D took reaction time data through a gamified version of the shortened 3-choice reaction time tests. Gamma distribution modes supported linear predictions for all three protocols, although Roll was the weakest. Roll (n = 14) was the least steep and was slightly asymmetric with a steeper positive slope than negative slope while Yaw (n = 14) was symmetric with an intermediate slope. Pitch (n = 14) was the steepest and most asymmetric, also with a steeper positive slope than negative slope. Unlike the other groups, Roll had the weakest linear prediction with large reduced-chi-squares, indicating a nonlinear relation. Especially for positive angles, the data appeared to curve downwards at larger angles, with a smaller reaction time for full rotation than for intermediate rotations up to 120°. Yaw and Pitch appeared mostly linear but also had large reduced-chi-squares. However, except for positive yaw, these values were reasonable given the small error bars. Positive yaw appeared linear excluding an increased reaction time at 90°.

For character roll, the Gamma distribution for English (n = 13) had a single mode below 1 while Thai (n = 13) and Chinese (n = 13) had modes greater than 1, implying a weaker linear prediction. Likewise, the aggregated analysis for English resulted in a strong linear model that was steeper than Groups A and B, making it half as steep as face roll with a 35 ms difference between unrotated and fully rotated characters. The linear predictions for Thai and Chinese were parallel to the English prediction, but high reduced chi-squares implied nonlinear relationships. In both cases, the data appears to plateau or curve downwards at higher rotations.

### 2.3 Comparison Between Groups

Although each group took data independently with different experimental setups and protocols, all four groups achieved similar V-shape patterns for corresponding protocols in both intercept and slope (Figure 5). Hence, data from each group could be pooled and analyzed as a single combined group.

**Figure 5.**
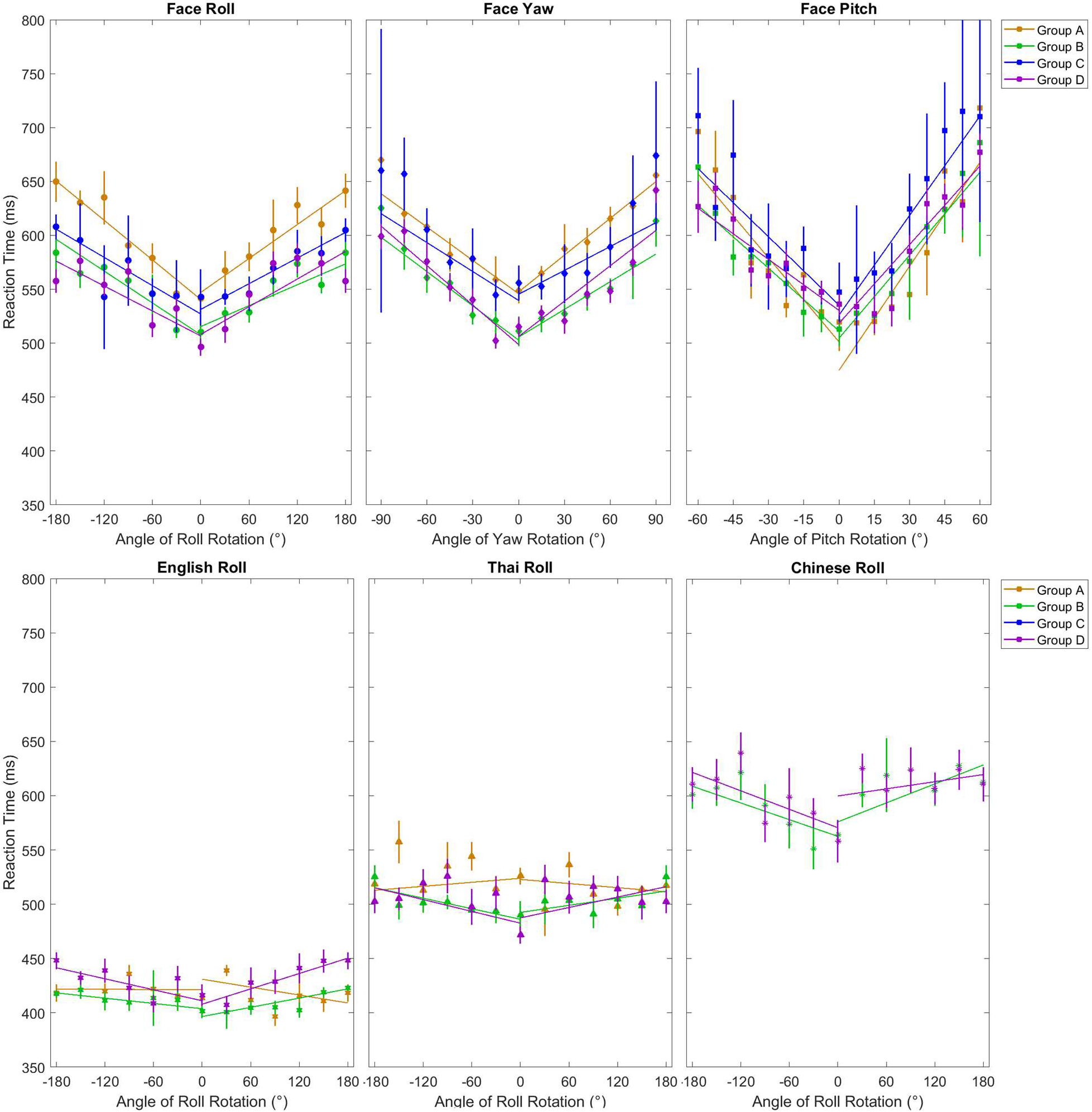
Protocol plots showing group aggregate fits on a common axis.

#### Comparison of Group Slopes

Slopes for Face Roll and Yaw between groups were consistently around 0.4 and 0.9 ms/°, respectively. On average, the V-shape for roll was symmetric while for yaw, it was slightly asymmetric with leftward yaw angles leading to longer reaction time shifts. Additionally, yaw was twice as steep as roll. In contrast, face pitch was consistently asymmetric between all groups, with upward rotations leading to longer reaction time shifts than downward rotations. All four protocols led to near-zero slopes for English and Thai Roll. Group A, which used different Thai characters, had an inverted v-pattern while groups B and D had normal but flat v-patterns. Chinese Roll resulted in steeper v-patterns that were symmetric for Group B but asymmetric for Group D.

**Figure 6.**
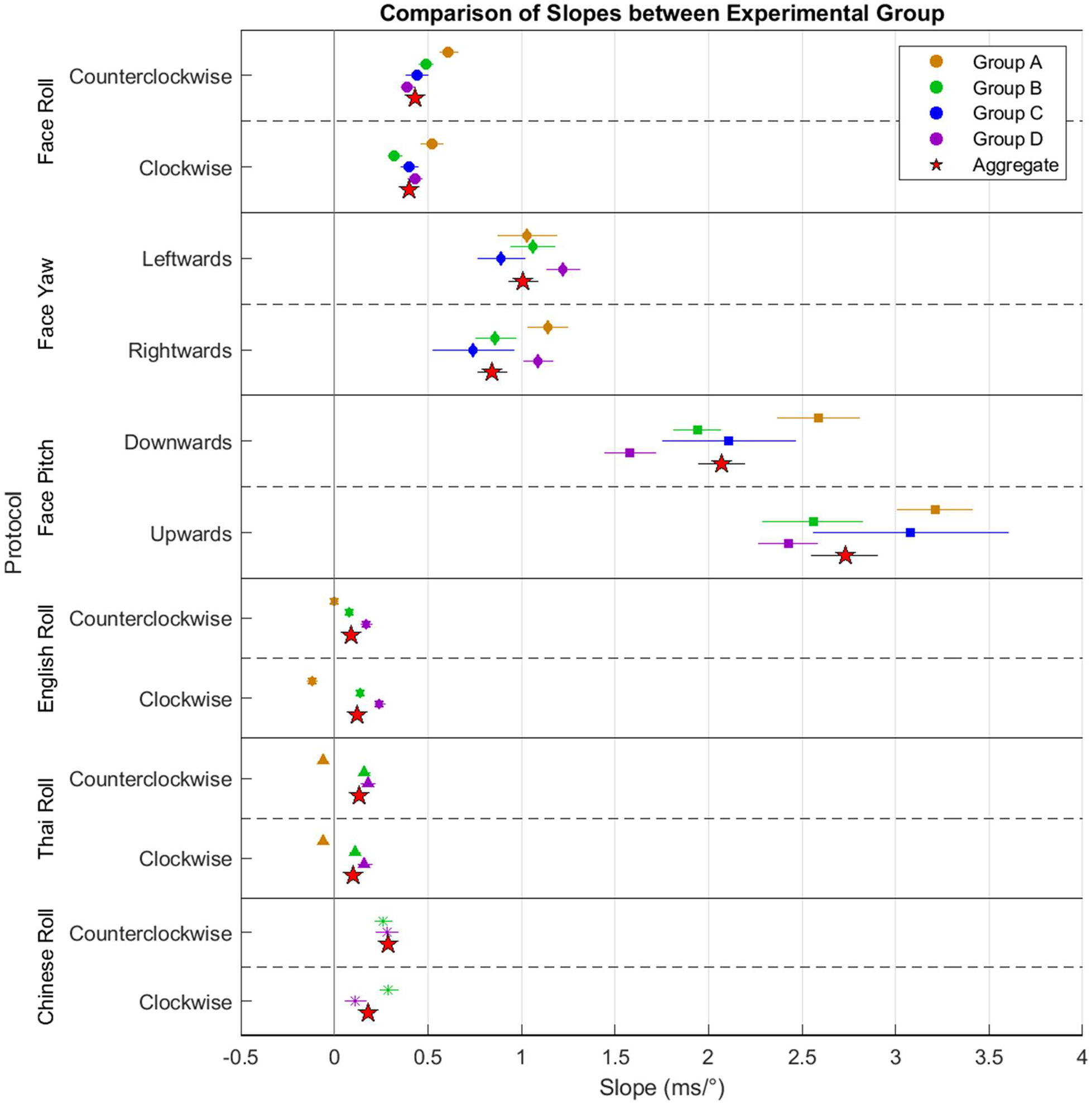
Comparison of influence of rotation on reaction times between Groups A-D for each protocol, represented by trendline slopes. Aggregate slopes (✩) were also included that combined all four participant groups.

**Figure 7.**
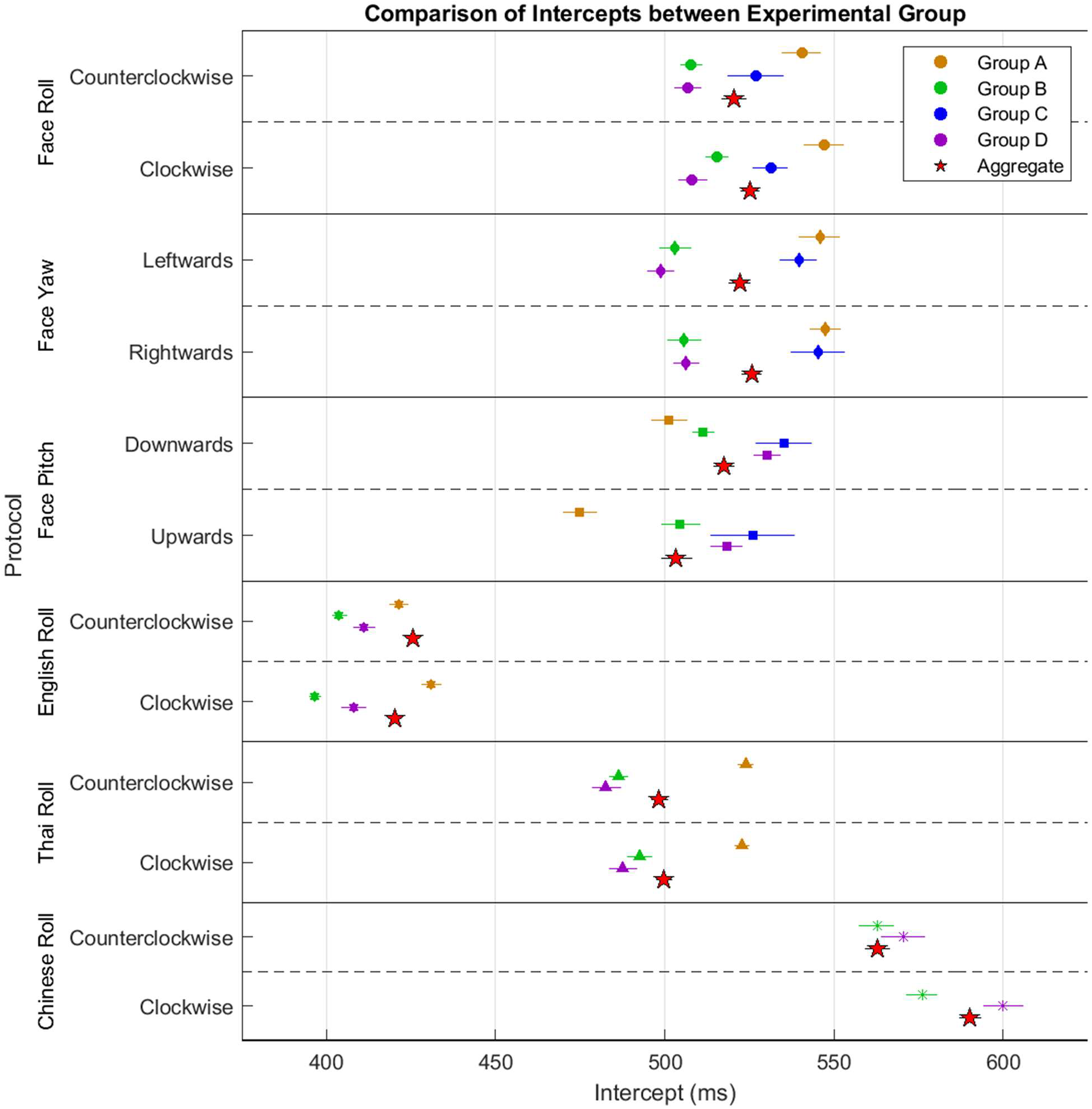
Comparison of reaction times for unrotated stimuli between Groups A-D for each protocol, represented by trendline intercepts. Aggregate intercepts (✩) were also included that combined all four participant groups.

#### Comparison of Group Intercepts

Within Groups B, C, and D, intercepts remained mostly constant between protocols, but Group C intercepts around 540 ms were notably higher than Groups B and D around 510 ms. Group A intercepts were also comparable to group C except for face pitch, which was faster than all other groups. Among character protocols, English had the fastest intercept time at 420 ms; Thai had an intermediate intercept time at 500 ms; and Chinese had the slowest intercept time at 575 ms.

## 3 Discussion

### 3.1 Linearity of Face Recognition in Three Rotational Axes

To support the new model of MePMoS and NHT, reaction time experiments were conducted for unfamiliar face and character rotation along the Roll, Yaw, and Pitch axes. Four independent groups, including two internal and two external groups, took data with variations between each group protocol and experimental environment. Nevertheless, agreement was found between corresponding protocols from each group. A positive linear correlation was found between reaction time and the degree of rotation of unfamiliar faces in all three axes, with the memorized front-facing, level, and upright faces having the fastest reaction time around 525 ms and larger angles of rotation having longer reaction times on average. These results support the NHT model, providing a way for the brain to transform memories of the faces by alpha wave phase shifts to overlap with incoming stimuli through time delays. Larger phase shifts caused by longer time delays could allow for recognition of larger rotation of the face in all three axes. The timescale of reaction times on the order of 100 ms also supports the involvement of alpha waves, with 525 ms for unrotated stimuli, a 75 ms shift for maximal Roll, a 100 ms shift for maximal Yaw, and a 125 ms shift for maximal Pitch.

As in previous studies, reaction times to rotations along the roll axis were generally faster than yaw and pitch (S. K. Favelle et al., 2011). The conversion to polar in the visual cortex V1 could aid in hastening roll rotation, making roll faster than the other two transformations. Roll rotation also occurs in 2D, so the log-polar ventral pathway is primarily needed to perform the rotation. In contrast, rotation along the pitch and yaw axes requires the brain to rotate a 3D memory of the stimulus and thus requires input from both the dorsal and ventral pathway (Todd, 2004). With involvement from both pathways, frame transformation between egocentric and object-centric references before application of the Yaw or Pitch rotation may contribute to the steeper reaction time shifts. Between Yaw and Pitch, Yaw was generally faster as previously observed (S. K. Favelle et al., 2007). By repeated practice or evolution, humans may have adapted to recognize yaw rotations more efficiently, likely due to the tendency to interact with others at a level height, leading to less exposure to faces at extreme pitch orientations than yaw orientations.

### 3.2 Unfamiliar Faces vs Characters

Unlike face rotation, character rotation was mostly flat with only slight changes in reaction time. English produced a strong positive linear correlation between reaction time and angle of rotation, but Thai and Chinese, both unfamiliar characters, had weaker linear correlations. Nevertheless, these results do not disagree with the proposed NHT model. For English and Thai, the characters assessed were arguably simpler than faces, as demonstrated by decreased central reaction times. Although unfamiliar characters took longer than familiar characters, the slopes were parallel, suggesting a shared method of rotation. Given the timescale of reaction time shifts at 10-15 ms, phase shifts in Gamma waves may encode rotation for simple shapes while alpha waves carry rotational information for more complex objects.

Chinese Roll data had a surprisingly higher central reaction time than even faces, most likely because of its unfamiliarity and a high density of key features such as corners and edges. It also had an intermediate slope between faces and simpler characters, revealing a possible relationship between the number of key features in a stimulus and whether the brain encodes visual information via Gamma waves, Alpha waves, or a mix of both. On the other hand, Chinese Roll appeared nonlinear which may reflect a divergence from holistic image processing to comparing the location of a few key features of the unfamiliar stimulus to memory, leading to faster but less accurate behavior. Hence, more experiments are required to better understand rotation of simple and complex characters, with an emphasis on the latter. In the current experiment, Chinese characters were selected by changing only a few radicals in each character set, leading to unilateral changes that could favor non-holistic differentiation based on feature detection. Introducing bilateral differences between characters in future experiments may allow for clearer characterization of the holistic approach used in more accurate language comprehension.

### 3.3 Systematic Error and Non-linear and Asymmetric Trends for Faces

There were some systematic errors present that were revealed by small differences between each group. For Group A, using different faces may have caused the inconsistencies in central reaction time between pitch and the other protocols. To reduce this error, the faces were standardized between protocols, which were taken in a randomized order. As a result, participants may have become more familiar with the faces for the second and third protocol, leading to reduced yet consistent central reaction times between each face protocol for Groups B-D.

The large variation in participant times for External Group C and the other test groups may have been caused by varying levels of motivation. Groups A and B consisted of highly motivated and knowledgeable research members who partook in designing the experiment. Consistent motivation for Group D was encouraged through added game-like components, including a new positive reinforcement system and score timer. On the contrary, Group C consisted of unknowledgeable participants, some of whom may not have been responding as quickly as possible while others did respond quickly. This may have increased variation in the slopes of the reaction time vs. angle of rotation plots, leading to larger error bars.

Especially for Group D, Roll rotations above 120 degrees were nonlinear. While there is a possibility that at such angles, the faces were still processed holistically yet very insufficiently (Richler et al., 2011), this finding is better explained by a qualitative shift in performance stemming from the loss of coherence between different facial features, which causes facial recognition to depend on some other non-holistic mode of processing. Upon reaching larger rotations, faces may be processed as objects, which suggests that faces may be processed less as a whole but by faster recognition of a few key features (Rossion, 2009). As the most motivated experimental group, Group D participants may have adopted this more “active” recognition to score more points when “passive” holistic processing took too long. Nevertheless, in daily life, it is likely that “passive” holistic processing through phase shifts of alpha waves is more prevalent due to its higher accuracy.

For Group D, Pitch had a significantly decreased reaction time at 22.5 degrees. This could result from different heights of participants to which the headrests were adjusted to allow comfortable data-taking. On average, participants may have been slightly above eye-level with the center of the screen, leading to the central reaction time when the stimuli faces were looking slightly upwards. However, given the distance between participants and the screen, this would only contribute to a small shift in the central angle. A more plausible cause could be the difference in overhead and ambient lighting between the professional and remote data-taking, which could influence perception of the faces (Favelle et al., 2017).

Yaw was slightly asymmetric while Pitch was highly asymmetric. For Yaw, this could represent systematic error from imperfectly symmetrical database faces, resulting in easier feature detection on one side of the face compared to the other. Likewise, differential lighting could also lead to differences in left and right face perception. On the contrary, asymmetry in Pitch can be explained from more exposure and practice to recognizing and processing more features from the top of other people’s heads than their chins.

### 3.4 Further Studies

In the current experiment, participants reacted fastest to the unrotated faces that they memorized. However, it is not known whether the brain automatically stores memorized faces at 0° or at the most practiced rotation. If the latter holds true, the observed V-pattern may translate horizontally to the angle at which training occurred. On the other hand, it is possible that the brain is biologically hardwired to automatically memorize novel faces as upright, front-facing, and level to facilitate daily social interactions.

Future experiments should also combine multiple transformations, such as multiple rotational axes or rotation and scaling. The transformations may simply add linearly or have cooperative or opposing effects, especially Yaw and Pitch rotations, which require similar steps to perform. The transformations may also contribute asymmetrically or even be independent where the final reaction time is determined by the alpha phase shift that normally takes more time. Finally, reaction time experiments could be used to better understand encoding of transformation of alphabets and words, which may yield similar results to faces. This could be due the transformations being the combination of multiple familiar Gamma-encoded characters, paralleling facial features in facial recognition.

This research supports a novel model of top-down human recognition of unfamiliar faces and the nuanced neurological pathway it adopts. We have provided evidence that visual processing involves the conversion of 7 degrees of transformation of images to time differences via a top-down pathway and the involvement of alpha and gamma waves phase shifts in encoding rotation. Top-down processing may better explain the incredible efficiency of stimulus recognition over bottom-up processing and how humans evolved to become intelligent beings that could recognize other individuals and comprehend language.

## 4 Materials and Methods

### 4.1 Participants and Equipment

Four groups of undergraduate students at the University of California, Los Angeles (UCLA), all of which had normal or corrected-to-normal vision, participated in the data-taking process. Due to the difficulties of recruiting participants during the COVID-19 pandemic, lab members from the UCLA Elegant Mind Club who designed the experiments also took data as internal participants. Groups A (n =13) and B (n = 15) consisted of internal participants involved with designing the experiment, respectively, who took data using personal computers and monitors of various sizes (Figure 8 Left). Participants created headrests from books or other household objects. Group C (n = 13) and D (n = 14) consisted of external participants, respectively, who took data in a professionally controlled laboratory setting using a 50-inch television, Sony headphones, and a custom-built headrest (Figure 8 Right). Participants for Group C were members of the Elegant Mind Club but were uninformed of the experiment and hypotheses while participants for Group D were completely external and recruited from lower division Physics lab classes. These groups’ data were assessed separately for consistency, then combined in a combined analysis as shown in the Results. Approval was given by the UCLA Institutional Review Board (IRB#19-001472-AM-00003) to conduct shorter experiments with external participants. All participants provided consent before completing the study and were debriefed afterwards.

**Figure 8.**
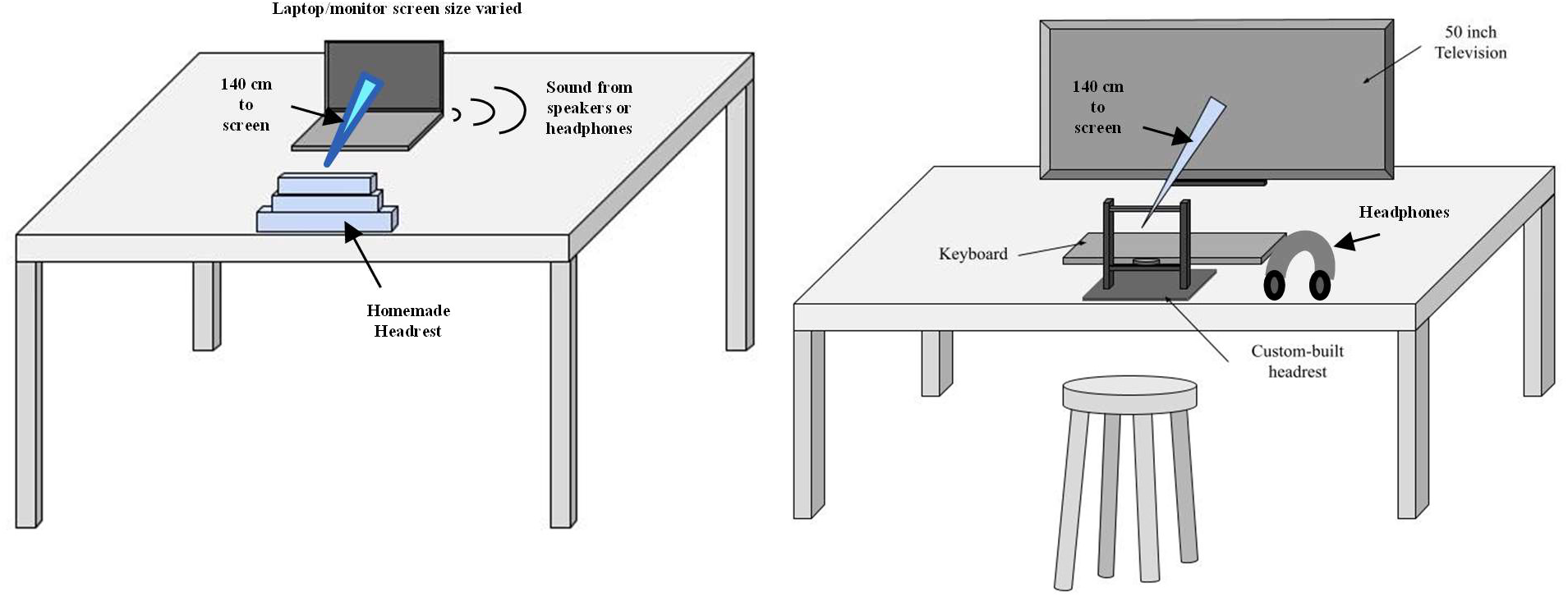
Drawing of experimental setups. (Left) Remote participants used makeshift headrests from household items to position their head 50 cm from their laptops or monitors with a keyboard within arm’s reach. (Right) Local participants took data in a controlled setting with a custom headrest 140 cm from a 50’’ TV and a keyboard within arm’s reach.

### 4.2 Display Calibration and Reaction-Time Consistency

To ensure consistency across devices, remote participants ran a size and time calibration script to control for figure display size and account for monitor delay parameters for the subsequent protocols. The size calibration determined the screen’s length to pixel conversion factor by prompting participants to resize vertical bars, letters, and images to various sizes using a ruler. The time calibration determined the input lag by showing a Gabor pattern on screen through PsychoPy and prompting users to press a key while recording themselves with a high-speed camera. A plot of PsychoPy times vs. actual times from the recording was generated to determine the time delay at the y-intercept.

### 4.3 Stimuli Selection

#### Unfamiliar Faces

Each face rotation experiment involved 9 emotionally-neutral adult face models, all of which were Caucasian and female, that were selected from the Stirling/ESRC 3D Face Database and exclusively grouped into sets of three based on similar age, face texture, skin tone, and head shape (*Stirling ESRC 3D Face Database*, 2021). Faces with less readily identifiable skin texture (freckles, moles, wrinkles) were preferred for experiments. Images were converted into grayscale to minimize the effects of differences in skin tone and blemishes and shown on a gray background. The models were rendered in Blender, an opensource 3D graphics software, with flat lighting and Gouraud Shading. Renders of the roll protocol were captured in 30-degree intervals from −150 to 180 degrees; the yaw protocol was captured in 15-degree intervals from −90 to 90 degrees; and the pitch protocol was captured in 7.5-degree intervals from −60 to 60 degrees.

#### Repeating faces and Randomization

For group A, the roll, yaw, and pitch protocols each had a different set of faces. For groups B-D, the protocols featured the same three sets of faces to minimize the systematic effect of using different faces. To negate the effects of the participants learning and familiarizing with the faces throughout the three protocols, participants were randomly assigned one of six permutations to run the face protocols. Stimuli images can be found in the repository linked in the Notes about Data and Authors section.

#### Character Selection

This study examined the effect of roll rotation for characters from three languages: English (familiar), Thai (unfamiliar), and Chinese (complex and unfamiliar). The Thai and Chinese protocols consisted of nine characters exclusively grouped into sets of three based on similar shapes and motifs such as curves and edges. All characters appeared white on a gray background in 30-degree intervals of roll rotation from - 150 to 180 degrees. For group A, B, and C, the English letters were a single set of uppercase EPB while group D used three sets of uppercase letters: EPF, OQG, and XYK. Group A used the different Thai characters than groups B-D. Group A did not record Chinese character data while groups B-D used the same set of 9 Chinese characters. Stimuli images can be found in the repository linked in the Notes about Data and Authors section.

### 4.4 Experimental Procedures

All experimental procedures followed the same general approach. In the learning period, participants were shown the unrotated stimuli for a duration of time after visual instructions to press a specific key whenever the stimulus was shown on screen. In the practice period, participants were assessed with unrotated stimuli without recording accuracy or reaction times while in the testing period, participants were tested with rotated stimuli, and accuracy and reaction times were recorded. Each trial began with a white cross flash at the center of the screen followed by a random pre-trial delay seconds before the stimulus appeared at the center of the screen (Figure 9). After pressing a key, the participants received feedback on whether they answered correctly or incorrectly. If participants responded incorrectly during testing periods, the missed trial was repeated later at the end of the set so that all angles were evaluated equally. This routine was repeated for each of the three sets of three stimuli with a 20 second break between each set. All faces and characters were displayed at an 8-degree pitch eccentricity from the eye’s opening angle. Participants from groups C and D were shown a tutorial video before each data taking session. Differences between each group are explained below and summarized in Table 4.

**Figure 9.**
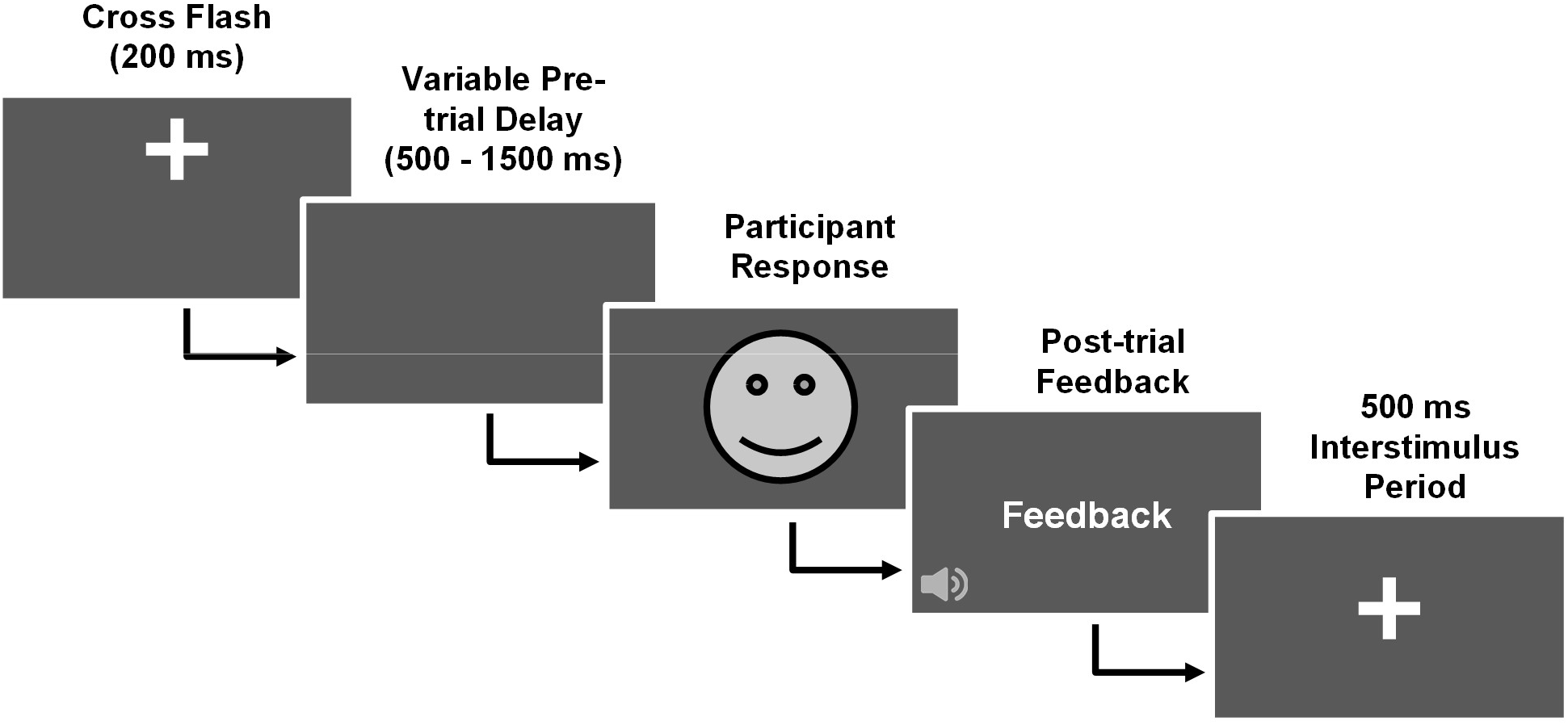
Flowchart showing steps for each trial. All faces and characters trials followed the same general steps, but there were intergroup differences in how participants responded and the audio/visual post-trial feedback

**Table 4.** Descriptions of four data taking samples. (*) As protocols were completed separately in a less controlled fashion, some internal participants did not complete all protocols. External participants also completed protocols over two days, so some only completed half of the protocols.

| Group Letter | Source of Participants | Max n* | Equipment | Experimental Design |
| --- | --- | --- | --- | --- |
| A | Internal members from Club | 14 | Laptops and various-sized monitors | Attractor vs. Distractor Test |
| B | Internal members from Club | 15 | Laptops and various-sized monitors | Full 3-choice Reaction Test |
| C | Semi-External members from Club | 13 | 50" TV, headphones, and headrest | Shortened 3-choice Reaction Test |
| D | External participants from Physics labs | 14 | 50" TV, headphones, and headrest | Shortened 3-choice Reaction Game |

#### Group A: Attractor vs. Distractor Test

In the learning period, only the attractor face or character was shown on screen for 30 seconds and mapped to the letter ‘v’. Participants were instructed to press ‘n’ when the unshown distractor faces and characters appeared. The practice periods consisted of 35 randomized trials with an equal probability of showing attractor and distractor stimuli, meaning attractors appeared twice as often as each individual distractor, and the testing periods consisted of 50 randomized trials. Halfway through the testing period, participants were given a shorter practice period with 15 trials. A solid A4 tone played for 0.5 seconds when participants pressed the wrong button. No sound played for a correct response.

#### Group B: Full 3-Choice Reaction Test

In the learning period, all three faces or characters were mapped to corresponding keys (v, b, or n). Participants were instructed to press a key before and after the corresponding stimulus appeared for 10 seconds. The learning period was repeated for a total of 20 seconds for each face. The practice periods consisted of 12 randomized trials with an equal probability of each stimulus appearing. For the full-length procedure, the testing periods consisted of 50 randomized trials with a shorter 6-trial practice period at the halfway mark. For the shortened procedure, training and practice periods were identical, but the testing periods consisted of 32 trials without an interim practice period. The same audio feedback was used as in the attractor-distractor test.

#### Group C: Shortened 3-Choice Reaction Test

Training and practice periods were identical to the full 3-choice reaction test. The testing periods were shortened to 32 trials per set without an interim practice period. Additionally, all three protocols were taken within one hour in a preset random order with a 1–4-minute break between each protocol. The audio feedback was used as in the attractor-distractor test.

#### Group D: Shortened 3-Choice Reaction Game

Gaming features were added to the training, practice, and testing periods of the shortened 3-choice reaction test. The instructions were modified to be more game-like with textual changes and animations for a short demonstration of the current rotation being assessed, a scoreboard showing 10 high scores, and a level screen. The scoring system was created with correct responses earning between 400 and 1000 points depending on response speed. A green message showed the score increase and a major triad played beginning with an A4 root that increased with consecutive correct answers. Incorrect responses resulted in a loss of up to 400 points with faster responses having a smaller decrease. After 1.2 seconds, the trial would time out, resulting in an automatic 400 score decrease. A red message showed the score decrease and an A4 tone followed by an F4 provided a defeat sound-effect.

### 4.5 Statistical Analysis

Reaction time trial data were categorized by their stimulus heights, and further analysis was done within the data separated by each height.

#### Error Analysis

The resulting reaction times were averaged, and the error bars were plotted using the calculated standard error, taking the standard deviation of the eccentricity angle reaction time divided by the square root of the number of trials within the eccentricity angles. After reaction time data was categorized by angle and outliers were removed, the standard deviation and standard error were calculated, with the standard error being used in the plots.

#### Outlier Analysis

Outlier data were removed before averaging the RT at each eccentricity. Data points that fell below the 25th percentile minus 1.5 times the interquartile range or above the 75th percentile plus 1.5 times the interquartile range for reaction time data at each angle were removed. The resulting data points were then averaged and used in chi-square calculations.

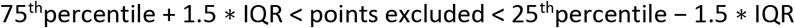

#### Hypothesis Testing

To ensure that an attention shift occurred during each choice RT block, response accuracy was analyzed using a one sample t-test for a mean of one, with three being the number of available choices for each protocol. The test was performed for each stimulus height and participant’s data was removed for a given height if the corresponding p-value was greater than 0.05.

#### Normalization of average RT

To better represent the relationship between RT and stimulus height in the aggregated analysis, we normalize each participant’s data to a global average for each protocol. We normalize the data by subtracting the appropriate participants’ mean performance from each observation, and then add the grand mean score to every observation. Let y be the i-th participants score in the j-th condition (i = 1,… N and j = 1,… M). Then define the normalized observations z-

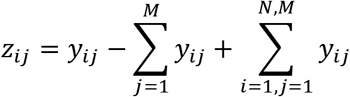

#### Line of best fit and correlation coefficient (χ^2^)

A best fit line was added using a least-squares linear regression for both positive and negative heights and their corresponding reaction times, with the memorized reference size being included in both positive and negative fits. The linear regression returned slope and intercept parameters as well as the correlation coefficient and a two-sided p-value in which the null hypothesis assumes a slope of zero. Additionally, the standard error of the fit was returned.

A best fit line of the form y = mx + b was calculated using chi-square minimization to find the best fit slope and intercept parameters for both positive and negative heights with each side including the reference height. From the chi-square minimization, error estimates for the slope and intercept parameters were computed by holding the other parameter constant at the best fit and calculating which parameter values would yield the minimum chi-square value + 1. This process was done for both slope and intercept parameters. The reduced chi-square was also calculated to determine goodness of fit by dividing the minimum chi-square value by the degrees of freedom. The correlation coefficient was also calculated using the NumPy library in Python.

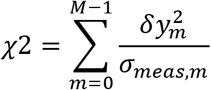

where 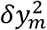 is defined as the mth residual 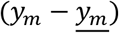 and *σ_meas,m_* is the standard error of the reaction time data for an angle m.

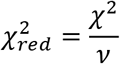

where *ν* is the degrees of freedom.

## Notes about Data and Authors

### Data availability

The experimental data, stimuli, and code to calibrate the monitors/TVs, analyze data, and run the experiments can be found at https://github.com/cm600286/EMC-Rotation-Station.

## Acknowledgements

This work was in part supported by the Dean’s Office of Life Science, Dean’s Office of Physical Science, Chair’s Office of the Department of Physics and Astronomy, and the Instructional Improvement grant by the Center for the Advancement of Teaching, all of which are at the University of California, Los Angeles. We would also like to extend our gratitude to the Arisaka Lab and the UCLA Physics & Astronomy for providing us working space and other resources. We are grateful for all those who have provided us with human face image/model repositories. Finally, we thank the banana for being a good model of holography.

## Author contributions

KA developed the concept and supervised the project. C.L, S.P., C.M., B.T., H.S., E.B., S.N, J.C., J.T, A.C., A.Z., D.W., N.C., E.L., G.K., E.O., F.W., A.Y., A.N., K.W., and P.W. wrote the manuscript. The research was conducted by two groups. In the “senior” group, N.C., B.T., E.M., E.L., S.N., F.W., A.Y., P.W., T.M., U.A., C.M., G.K., A.N., E.O., K.W., M.M., A.L., and L.S. designed and conducted initial experiments for experimental Group A. The “new” group, comprised of C.L., N.C., J.C., S.P., B.T., D.W., A.Z., J.T., A.C., E.B., and H.S., redesigned the experiments with the help of B.T and P.G. and conducted experiments as operators for groups B, C, and D. D.E. managed IRB approval and participant recruitment, and D.E., J.C., and new group members set up hardware for external data taking. B.T, P.W., T.M., and C.L created PsychoPy and MATLAB scripts. P.G. and N.C. created stimulus images for the senior group, and N.C., C.L., J.C. and A.Z., selected and created stimuli for the new group. N.C., B.T., E.L., S.N., E.O., U.A., and P.W. performed statistical analysis for the senior group, and C.L., S.P., J.T., A.C., and D.W. performed final statistical analysis and figure generation. N.C., E.L., S.N., F.W., and A.Y. coordinated the senior group process, and C.L., J.C., S.P., and D.W. coordinated the new group progress. K.A. and B.T. supervised the work, and K.A. and A.B. finalized the manuscript. All authors have looked at and approved the manuscript.

