## Supplementary Information for "Visual Perception of 3D Space and Shape in Time - Part IV - 3D Shape Recognition by 3D Rotation"

| Experiment Information | | | Negative Angles (Counterclockwise Roll, Leftward Yaw, Downward Pitch) | | | | | Positive Angles (Clockwise Roll, Rightward Yaw, Upward Pitch) | | | | |
|---|---|---|---|---|---|---|---|---|---|---|---|---|
| Protocol | Group | n | Slope (ms/°) | Intercept (ms) | R | $\chi^2$ | Reduced $\chi^2$ | Slope (ms/°) | Intercept (ms) | R | $\chi^2$ | Reduced $\chi^2$ |
| Face Roll | A | 14 | -0.61 ± 0.05 | 540 ± 6 | -0.97 | 1.07 | 0.21 | 0.52 ± 0.06 | 547 ± 6 | 0.95 | 2.54 | 0.51 |
| | B | 15 | -0.49 ± 0.04 | 508 ± 3 | -0.95 | 6.10 | 1.22 | 0.32 ± 0.04 | 515 ± 4 | 0.92 | 6.05 | 1.21 |
| | C | 13 | -0.44 ± 0.06 | 527 ± 8 | -0.81 | 1.15 | 0.23 | 0.40 ± 0.05 | 531 ± 5 | 0.96 | 0.78 | 0.16 |
| | D | 14 | -0.39 ± 0.04 | 507 ± 4 | -0.83 | 12.91 | 2.58 | 0.43 ± 0.04 | 508 ± 4 | 0.81 | 16.82 | 3.36 |
| | A/B/D | 43 | -0.47 ± 0.03 | 528 ± 3 | -0.98 | 3.73 | 0.75 | 0.43 ± 0.03 | 531 ± 3 | 0.92 | 10.46 | 2.09 |
| | Aggregate | 56 | -0.43 ± 0.03 | 521 ± 4 | -0.99 | 1.26 | 0.25 | 0.40 ± 0.03 | 525 ± 3 | 0.94 | 8.40 | 1.68 |
| Face Yaw | A | 14 | -1.03 ± 0.16 | 546 ± 6 | -0.96 | 1.12 | 0.22 | 1.14 ± 0.11 | 548 ± 5 | 0.99 | 0.37 | 0.07 |
| | B | 15 | -1.06 ± 0.12 | 503 ± 5 | -0.97 | 2.32 | 0.46 | 0.86 ± 0.11 | 506 ± 5 | 0.95 | 2.60 | 0.52 |
| | C | 13 | -0.89 ± 0.13 | 540 ± 6 | -0.94 | 4.11 | 0.82 | 0.74 ± 0.22 | 545 ± 8 | 0.90 | 1.98 | 0.40 |
| | D | 14 | -1.22 ± 0.09 | 499 ± 4 | -0.96 | 9.73 | 1.95 | 1.09 ± 0.08 | 506 ± 4 | 0.88 | 20.20 | 4.04 |
| | A/B/D | 43 | -1.05 ± 0.08 | 527 ± 3 | -0.98 | 2.93 | 0.59 | 0.98 ± 0.06 | 530 ± 3 | 0.96 | 8.89 | 1.78 |
| | Aggregate | 56 | -1.01 ± 0.08 | 522 ± 3 | -0.98 | 4.03 | 0.81 | 0.84 ± 0.08 | 526 ± 3 | 0.95 | 5.10 | 1.02 |
| Face Pitch | A | 12 | -2.59 ± 0.22 | 501 ± 5 | -0.94 | 12.57 | 1.80 | 3.21 ± 0.20 | 475 ± 5 | 0.92 | 17.34 | 2.48 |
| | B | 15 | -1.94 ± 0.13 | 511 ± 3 | -0.96 | 2.95 | 0.42 | 2.56 ± 0.27 | 505 ± 6 | 0.98 | 3.11 | 0.44 |
| | C | 13 | -2.11 ± 0.36 | 535 ± 8 | -0.87 | 4.26 | 0.61 | 3.08 ± 0.52 | 526 ± 12 | 0.96 | 2.27 | 0.32 |
| | D | 14 | -1.58 ± 0.14 | 530 ± 4 | -0.92 | 11.88 | 1.70 | 2.43 ± 0.16 | 518 ± 5 | 0.93 | 13.60 | 1.94 |
| | A/B/D | 41 | -1.99 ± 0.11 | 525 ± 3 | -0.97 | 5.67 | 0.81 | 2.66 ± 0.15 | 514 ± 4 | 0.96 | 17.22 | 2.46 |
| | Aggregate | 54 | -2.07 ± 0.13 | 517 ± 3 | -0.97 | 5.94 | 0.85 | 2.73 ± 0.18 | 503 ± 5 | 0.97 | 14.97 | 2.14 |

**Table S1** All parameters for unfamiliar face rotation. Aggregate values labeled A/B/D were also determined without Group C data.

#### Supplementary Information

| Experiment Information | | | Negative Angles (Counterclockwise) | | | | | Positive Angles (Clockwise) | | | | |
|---|---|---|---|---|---|---|---|---|---|---|---|---|
| Protocol | Group | n | Slope (ms/°) | Intercept (ms) | R | $\chi^2$ | Reduced $\chi^2$ | Slope (ms/°) | Intercept (ms) | R | $\chi^2$ | Reduced $\chi^2$ |
| English Roll | A | 9 | 0.00 ± 0.02 | 421 ± 3 | -0.24 | 4.63 | 0.93 | -0.12 ± 0.03 | 431 ± 3 | -0.23 | 20.03 | 4.01 |
| | B | 8 | -0.08 ± 0.02 | 404 ± 2 | -0.80 | 1.23 | 0.25 | 0.14 ± 0.01 | 397 ± 2 | 0.84 | 3.92 | 0.78 |
| | C | 0 | - | - | - | - | - | - | - | - | - | - |
| | D | 13 | -0.17 ± 0.03 | 411 ± 3 | -0.72 | 6.84 | 1.37 | 0.24 ± 0.03 | 408 ± 4 | 0.94 | 2.46 | 0.49 |
| | Aggregate | 30 | -0.09 ± 0.01 | 426 ± 2 | -0.88 | 1.29 | 0.26 | 0.12 ± 0.02 | 420 ± 2 | 0.89 | 3.17 | 0.63 |
| Thai Roll | A | 3 | 0.06 ± 0.02 | 524 ± 2 | -0.15 | 12.91 | 2.58 | -0.06 ± 0.02 | 523 ± 2 | -0.15 | 10.33 | 2.07 |
| | B | 9 | -0.16 ± 0.03 | 486 ± 3 | -0.81 | 2.50 | 0.50 | 0.11 ± 0.03 | 493 ± 4 | 0.64 | 4.68 | 0.94 |
| | C | 1 | - | - | - | - | - | - | - | - | - | - |
| | D | 13 | -0.18 ± 0.04 | 483 ± 4 | -0.46 | 10.52 | 2.10 | 0.16 ± 0.04 | 487 ± 4 | 0.27 | 14.52 | 2.90 |
| | Aggregate | 25 | -0.13 ± 0.02 | 498 ± 2 | -0.77 | 5.67 | 1.13 | 0.10 ± 0.02 | 500 ± 2 | 0.53 | 7.34 | 1.47 |
| Chinese Roll | A | 0 | - | - | - | - | - | - | - | - | - | - |
| | B | 9 | -0.26 ± 0.05 | 563 ± 5 | -0.83 | 3.34 | 0.67 | 0.29 ± 0.05 | 576 ± 5 | 0.67 | 9.13 | 1.83 |
| | C | 1 | - | - | - | - | - | - | - | - | - | - |
| | D | 13 | -0.28 ± 0.06 | 571 ± 6 | -0.73 | 6.49 | 1.30 | 0.11 ± 0.06 | 600 ± 6 | 0.52 | 8.90 | 1.78 |
| | Aggregate | 22 | -0.29 ± 0.03 | 563 ± 4 | -0.82 | 10.36 | 2.07 | 0.18 ± 0.03 | 590 ± 3 | 0.60 | 23.86 | 4.77 |

**Table S2** All parameters for English, Thai, and Chinese roll. Some samples were excluded for having too few participants

### Supplementary Information

#### Figure S1

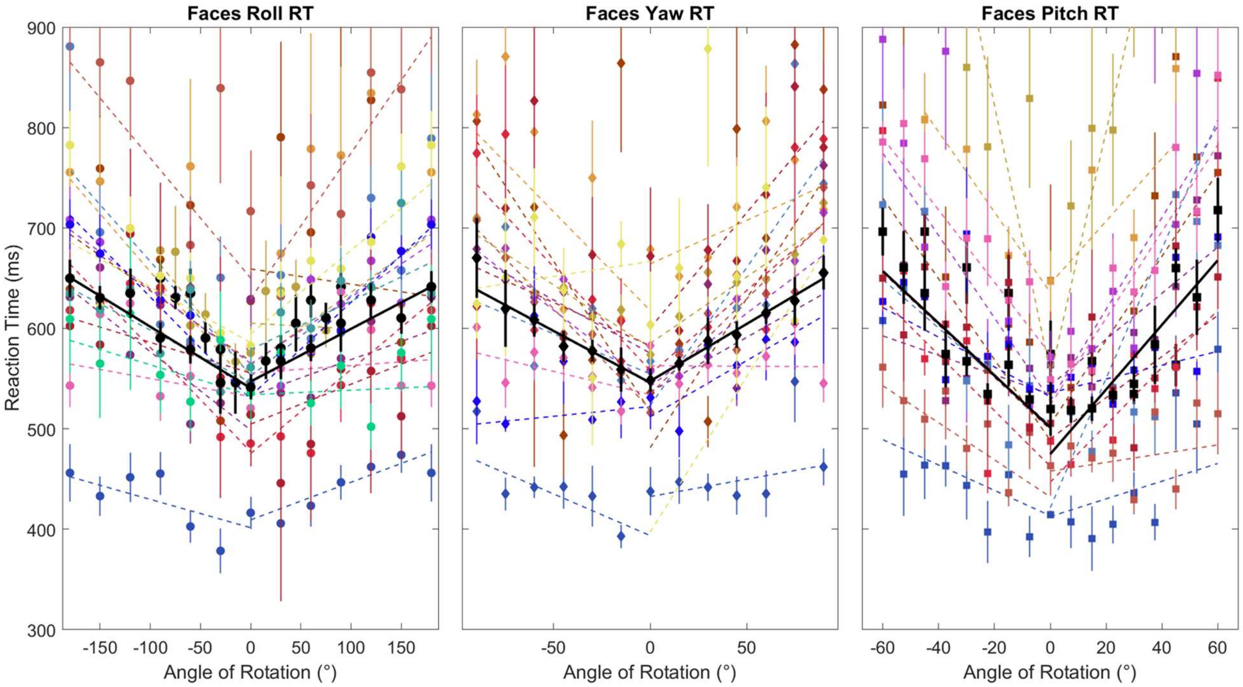


**Figure S1A** Group A raw reaction times for faces against roll, yaw, and pitch rotations. Error bars were calculated after a Cousineau normalization was applied.

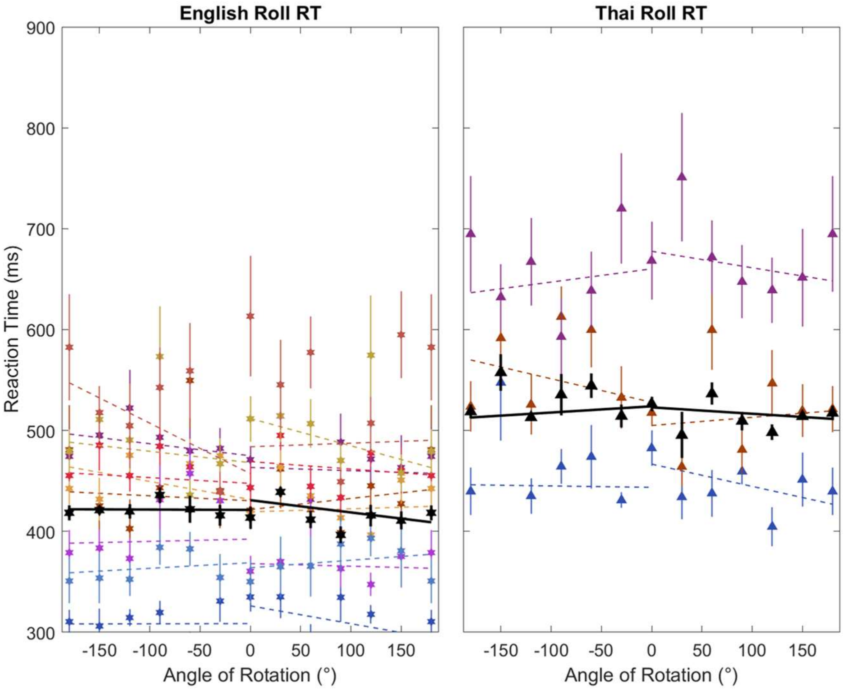


**Figure S1B** Group A raw reaction times for English and Thai against roll rotation. Error bars were calculated after a Cousineau normalization was applied.

### Supplementary Information

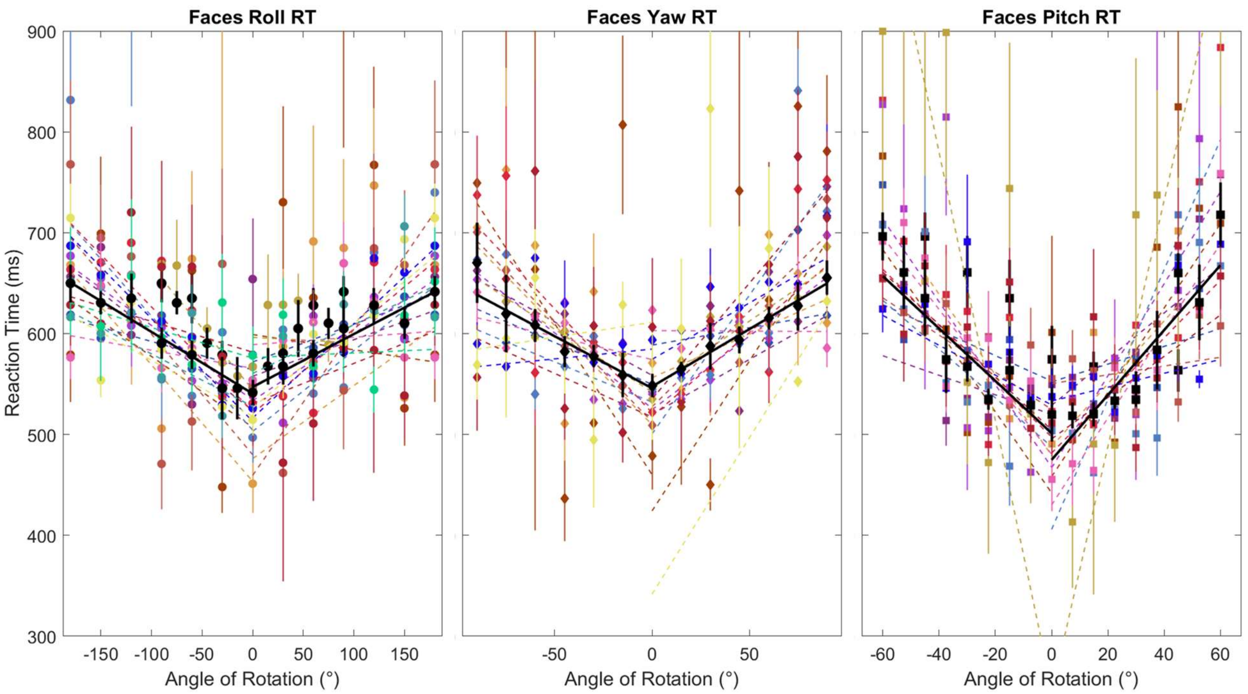


**Figure S1C** Group A normalized reaction times for faces against roll, yaw, and pitch rotations.

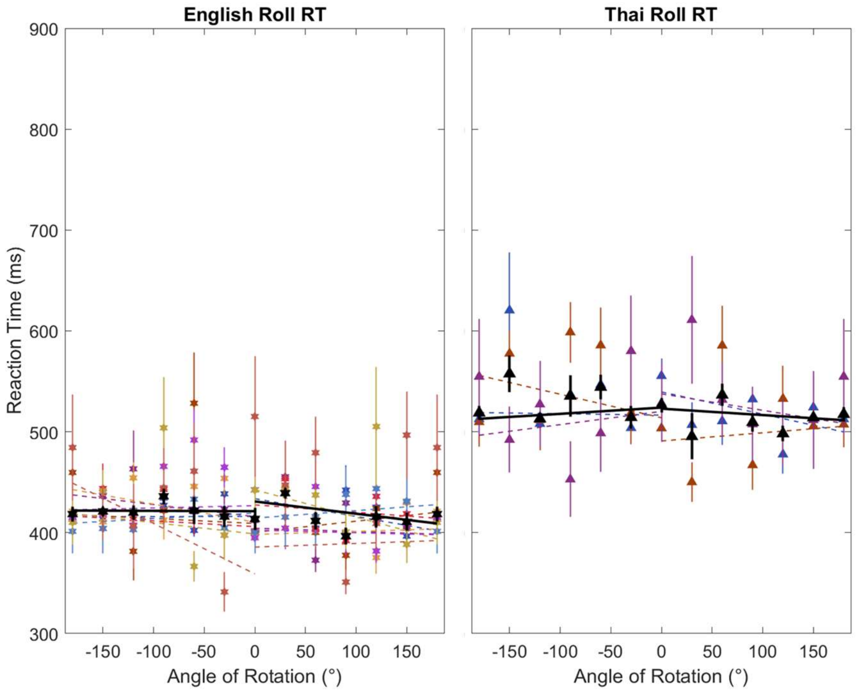


**Figure S1D** Group A normalized reaction times for English and Thai against roll rotation.

### Supplementary Information

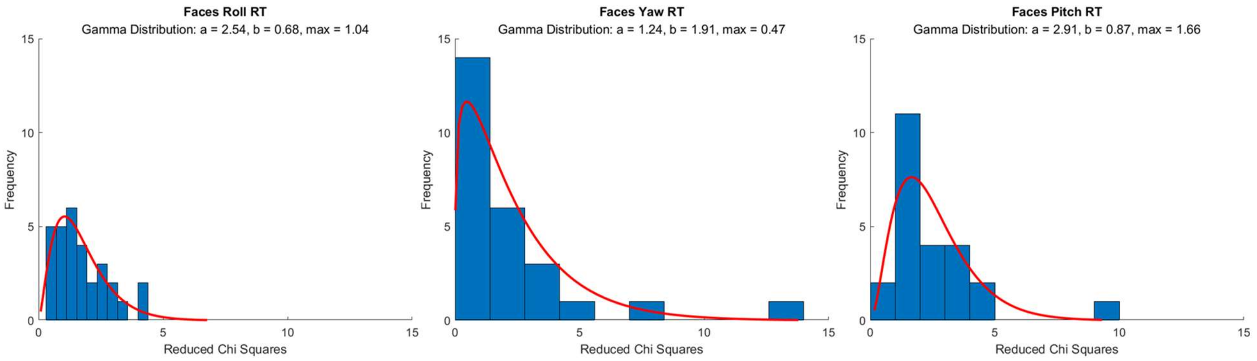


**Figure S1E** Group A distribution of reduced chi squares for individual participant face data. Reduced chi squares for linear fits of both positive and negative angles were analyzed together along with a gamma distribution curve. The data for each rotation direction was divided into 10 bins.

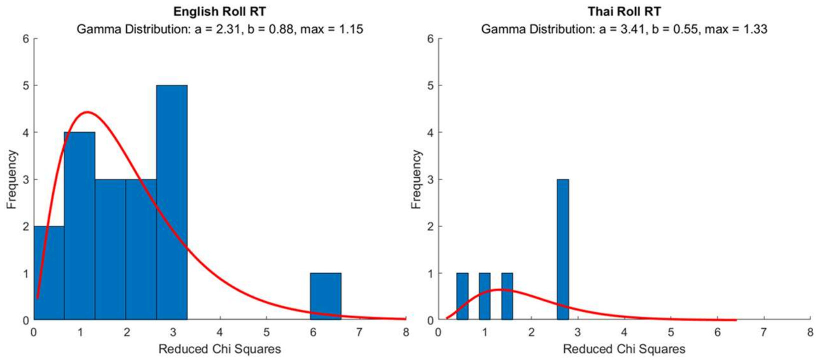


**Figure S1F** Group A distribution of reduced chi squares for individual participant character data. Reduced chi squares for linear fits of both positive and negative angles were analyzed together along with a gamma distribution curve. The data for each character type was divided into 10 bins.

### Supplementary Information

### Figure S2

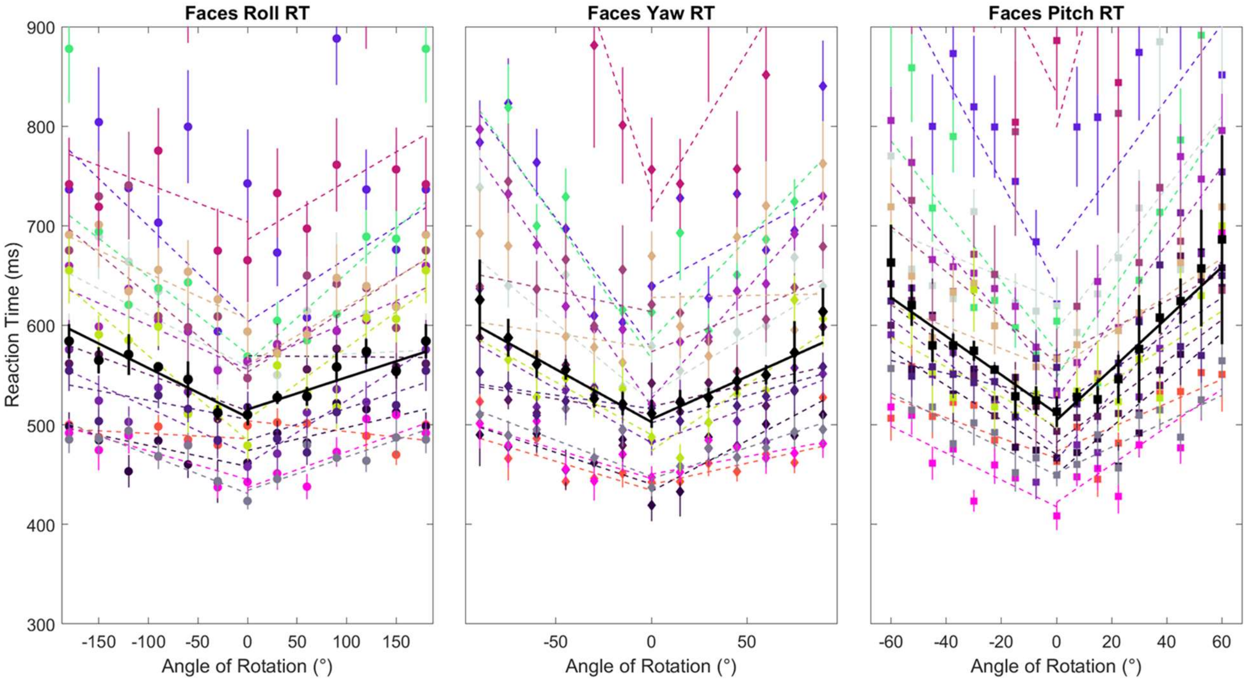


**Figure S2A** Group B raw reaction times for faces against roll, yaw, and pitch rotations. Error bars were calculated after a Cousineau normalization was applied.

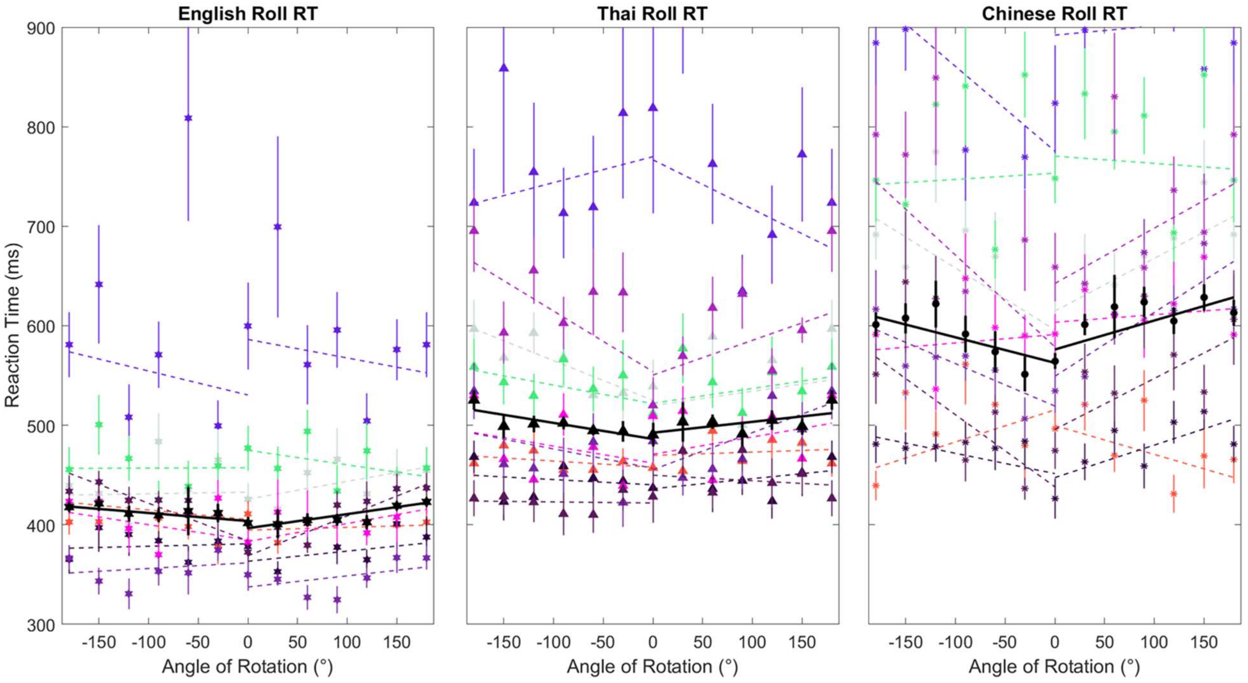


**Figure S2B** Group B raw reaction times for English, Thai, and Chinese characters against roll rotation. Error bars were calculated after a Cousineau normalization was applied.

### Supplementary Information

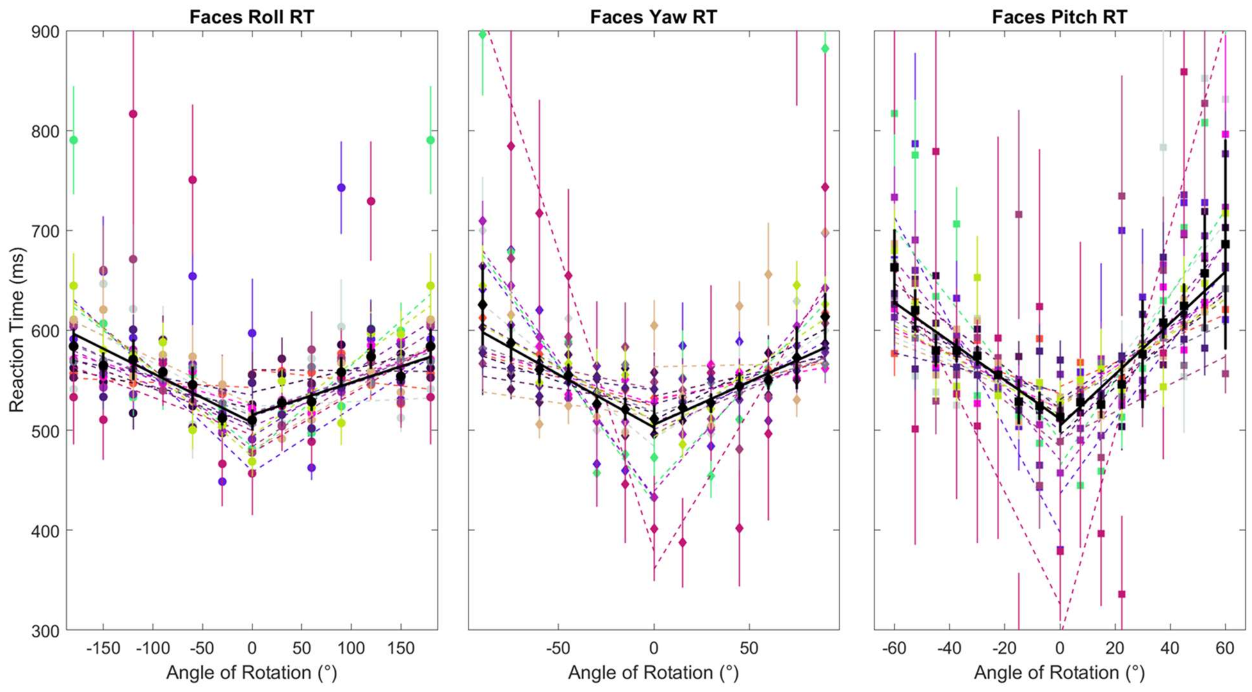


**Figure S2C** Group B normalized reaction times for faces against roll, yaw, and pitch angles.

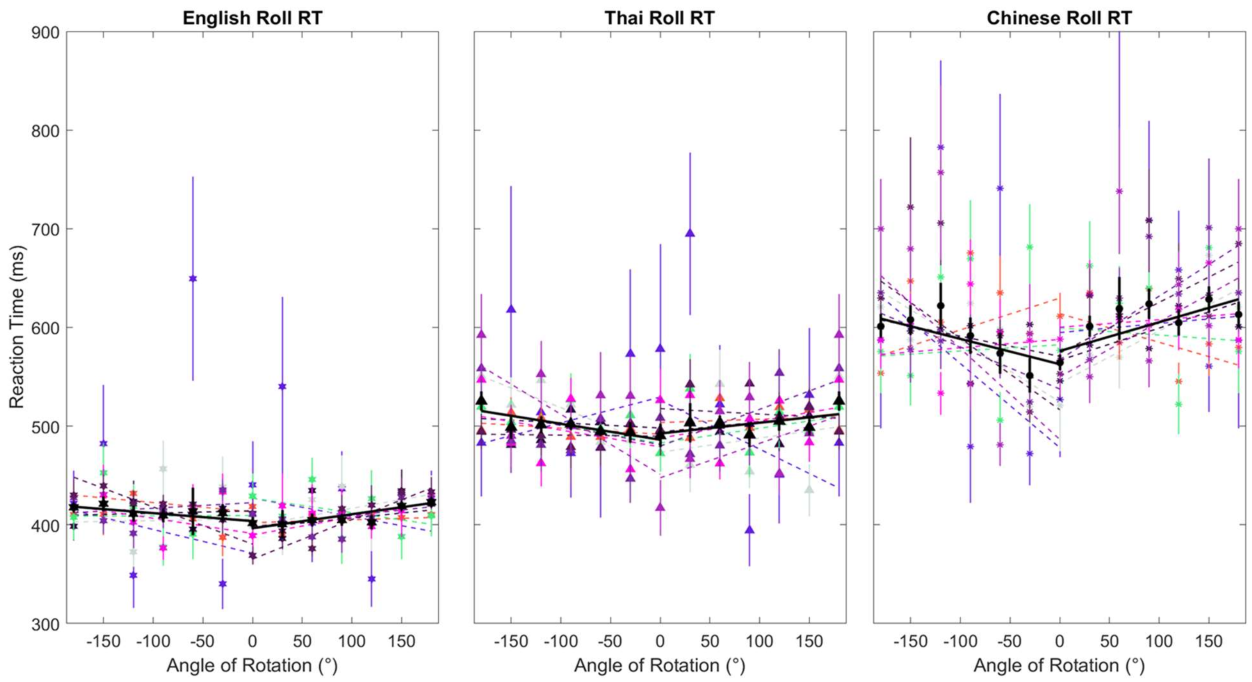


**Figure S2D** Group B normalized reaction times for English, Thai, and Chinese against roll angle.

### Supplementary Information

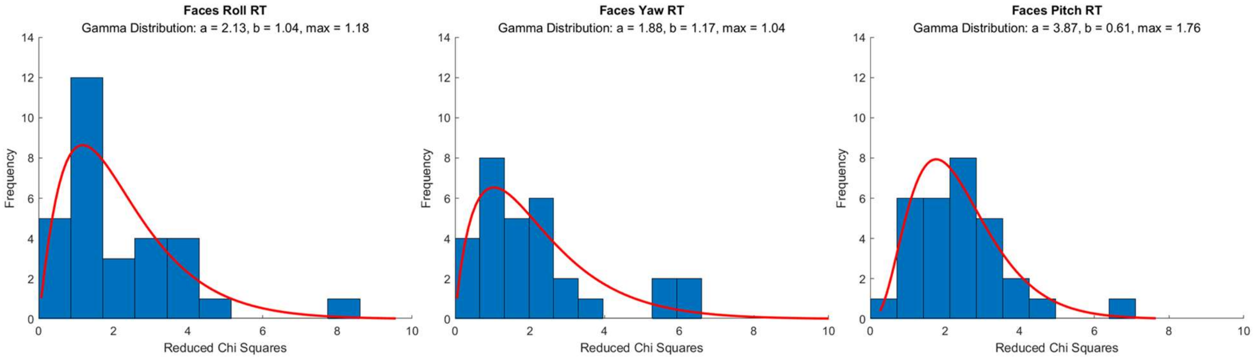


**Figure S2E** Group B distribution of reduced chi squares for individual participant face data. Reduced chi squares for linear fits of both positive and negative angles were analyzed together along with a gamma distribution curve. The data for each rotation direction was divided into 10 bins.

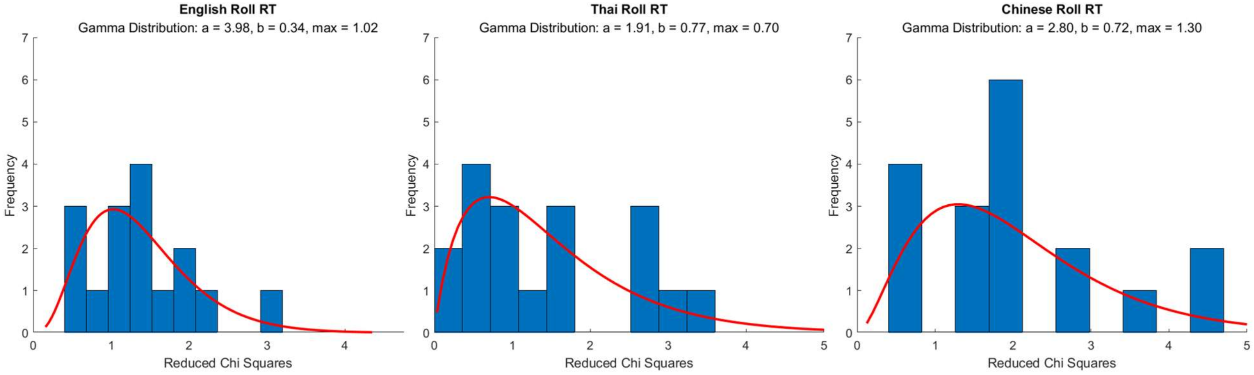


**Figure S2F** Group B distribution of reduced chi squares for individual participant character data. Reduced chi squares for linear fits of both positive and negative angles were analyzed together along with a gamma distribution curve. The data for each character type was divided into 10 bins.

### Supplementary Information

### Figure S3

Figures S6B, S6D, and S6F are omitted because Group C did not take character data.

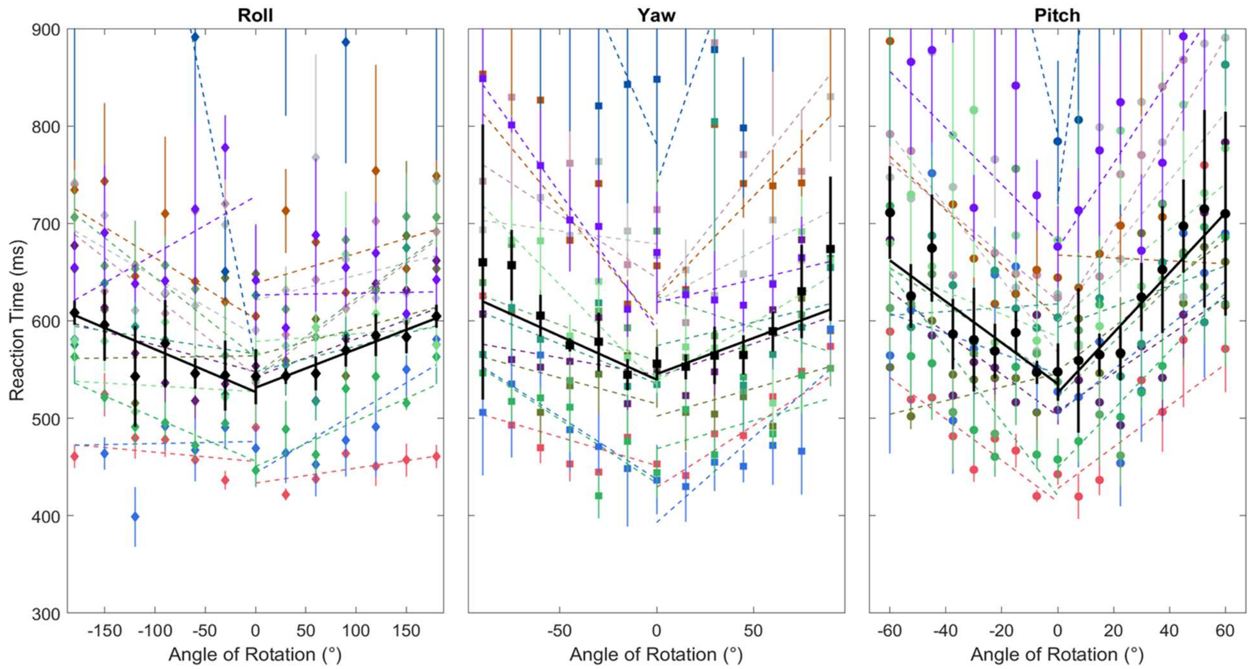


**Figure S3A** Group C raw reaction times for faces against roll, yaw, and pitch rotations. Error bars were calculated after a Cousineau normalization was applied.

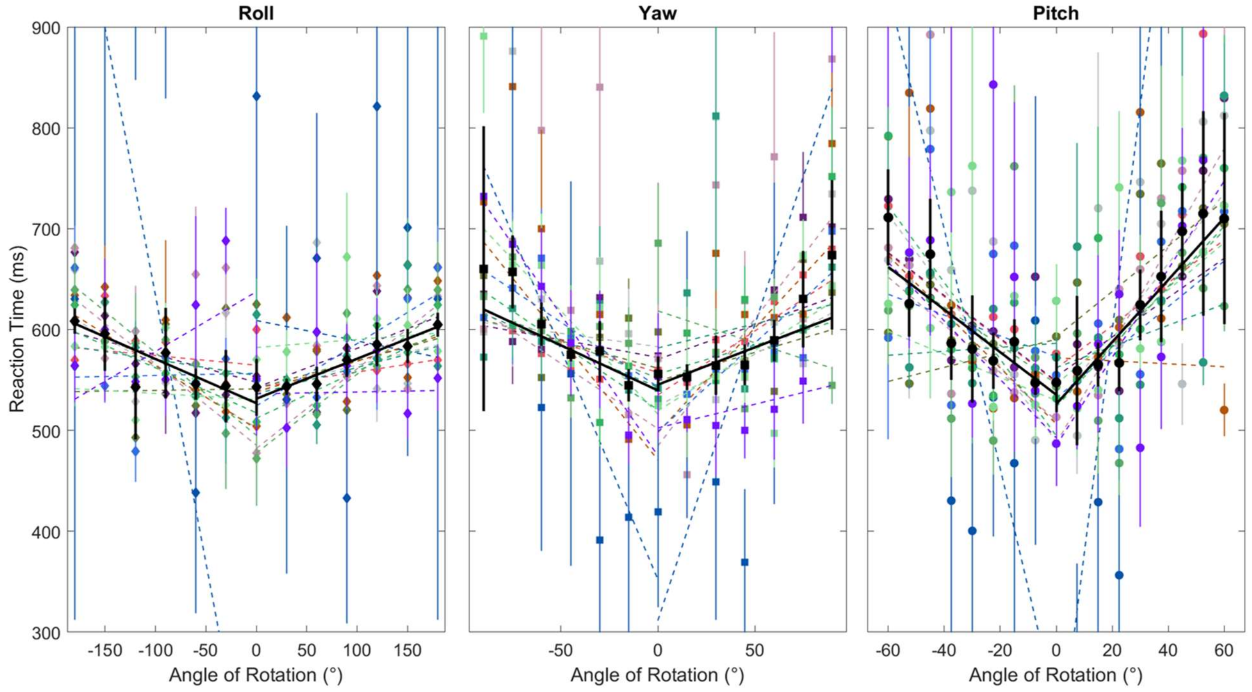


**Figure S3B** Group C normalized reaction times for faces against roll, yaw, and pitch rotations

### Supplementary Information

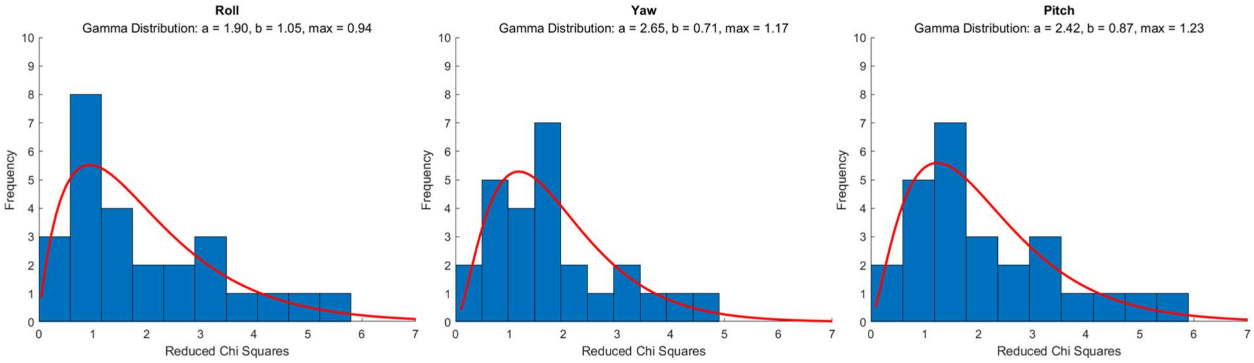


**Figure S3C** Group C distribution of reduced chi squares for individual participant faces data. Reduced chi squares for linear fits of both positive and negative angles were analyzed together along with a gamma distribution curve. The data for each rotation direction was divided into 10 bins.

### Supplementary Information

#### Figure S4

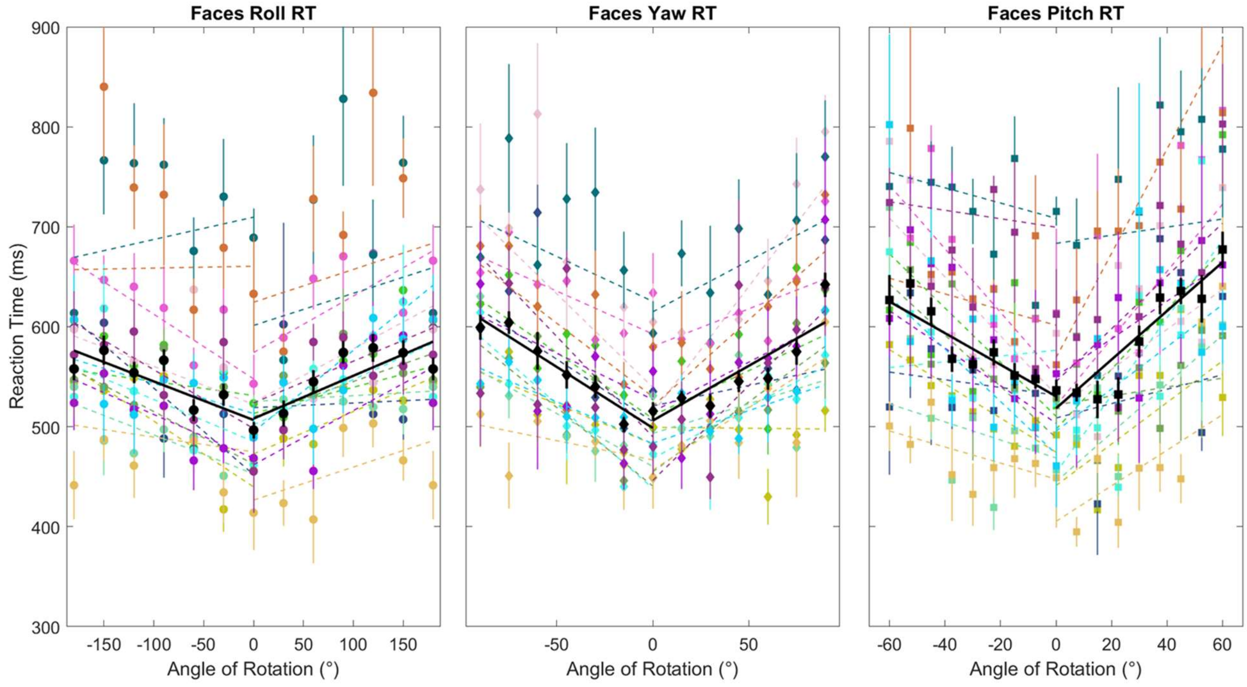


**Figure S4A** Group D raw reaction times for faces against roll, yaw, and pitch rotations. Error bars were calculated after a cousineau normalization was applied.

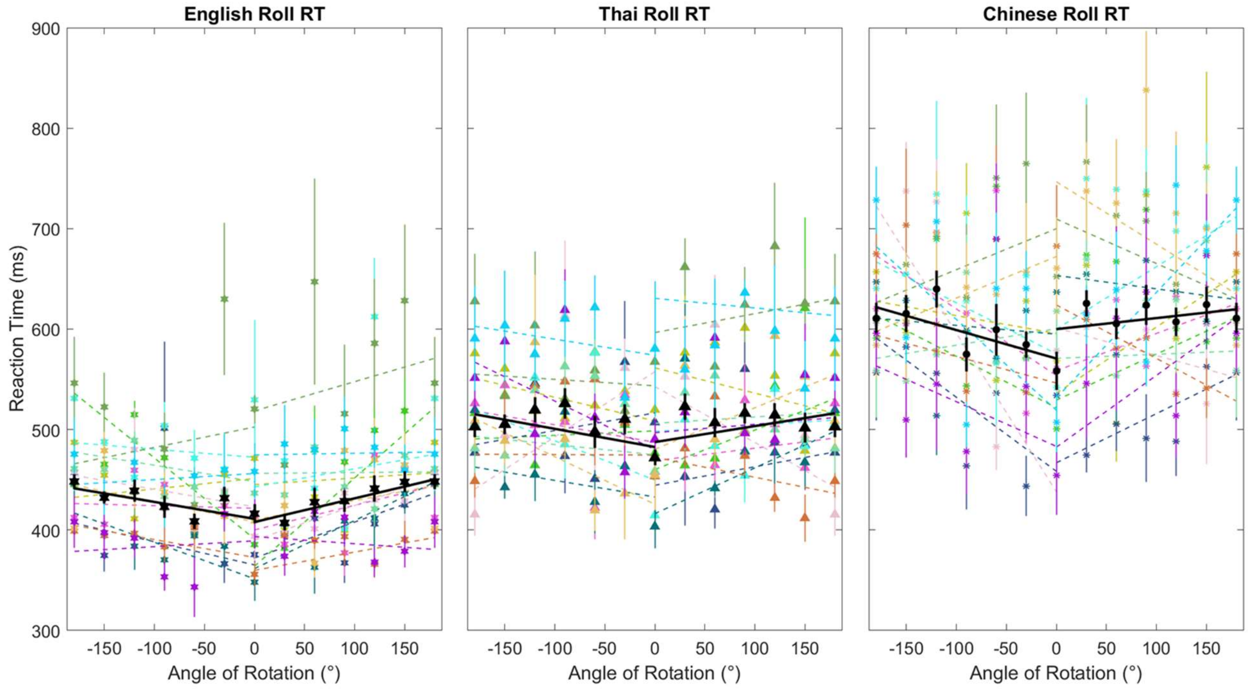


**Figure S4B** Group D raw reaction times for English, Thai, and Chinese characters against roll rotation. Error bars were calculated after a cousineau normalization was applied.

### Supplementary Information

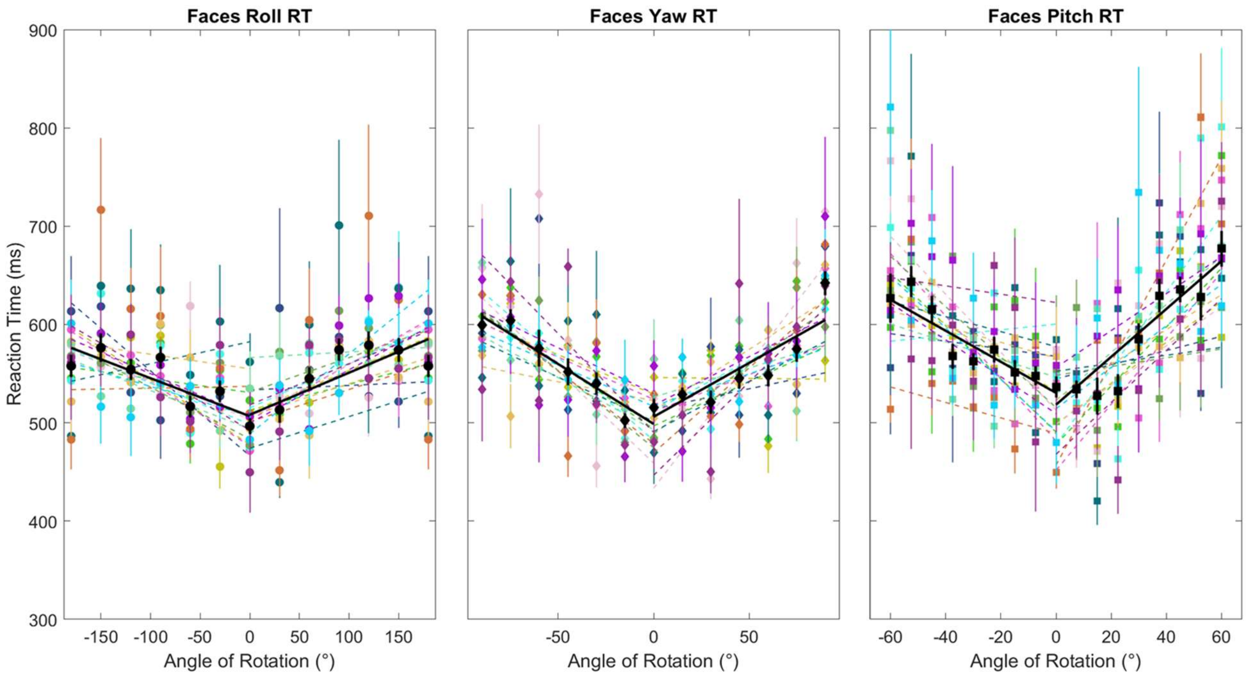


**Figure S4C** Group D normalized reaction times for faces against roll, yaw, and pitch rotations.

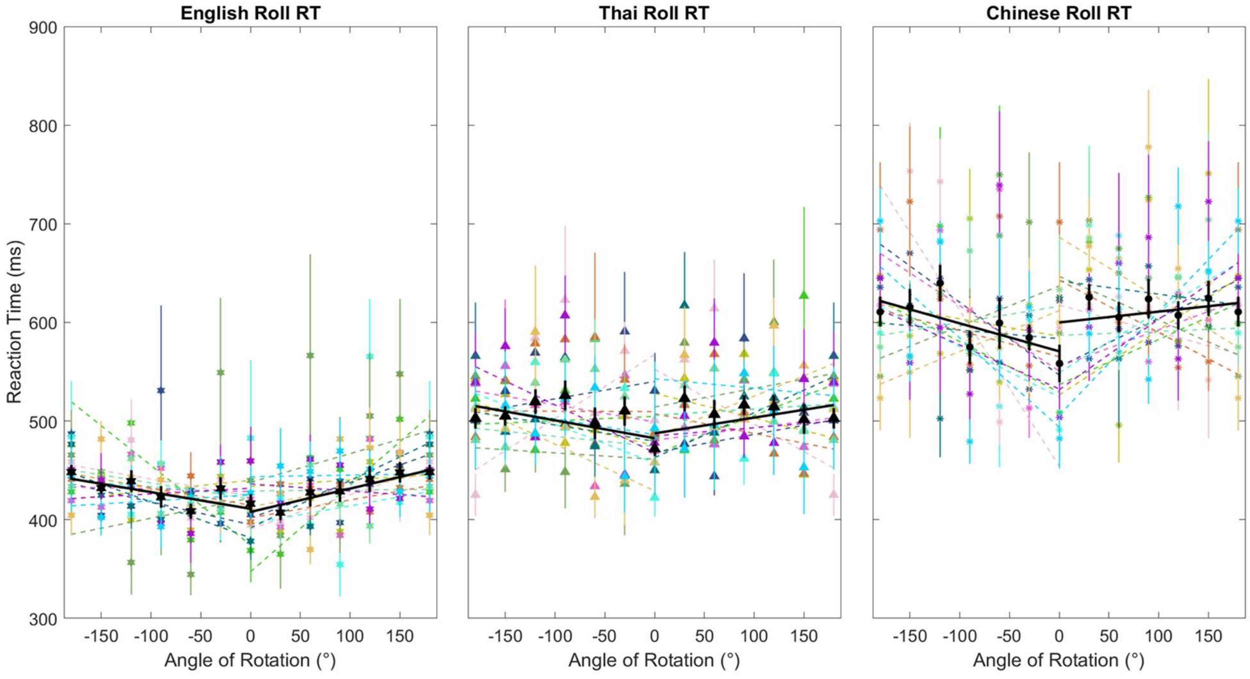


**Figure S4D** Group D normalized reaction times for English, Thai, and Chinese characters against roll rotation.

### Supplementary Information

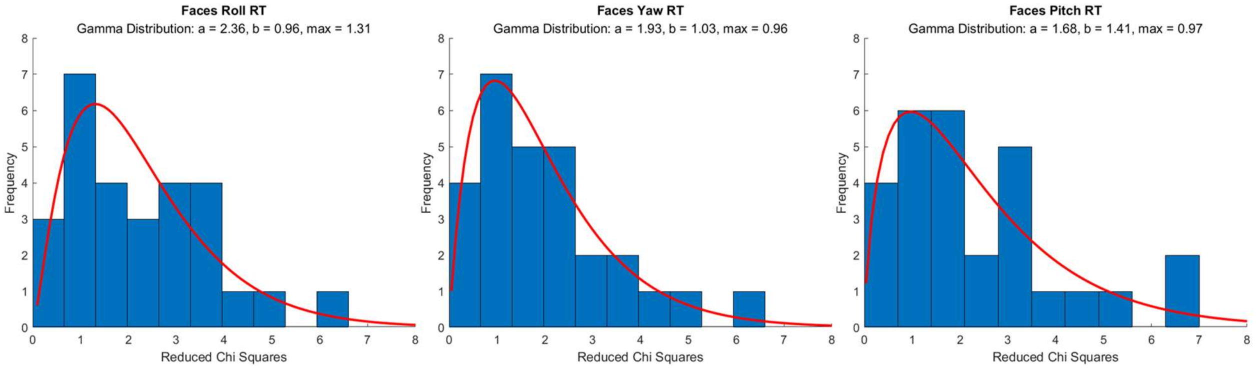


**Figure S4E** Group D distribution of reduced chi squares for individual participant faces data. Reduced chi squares for linear fits of both positive and negative angles were analyzed together along with a gamma distribution curve. The data for each rotation direction was divided into 10 bins.

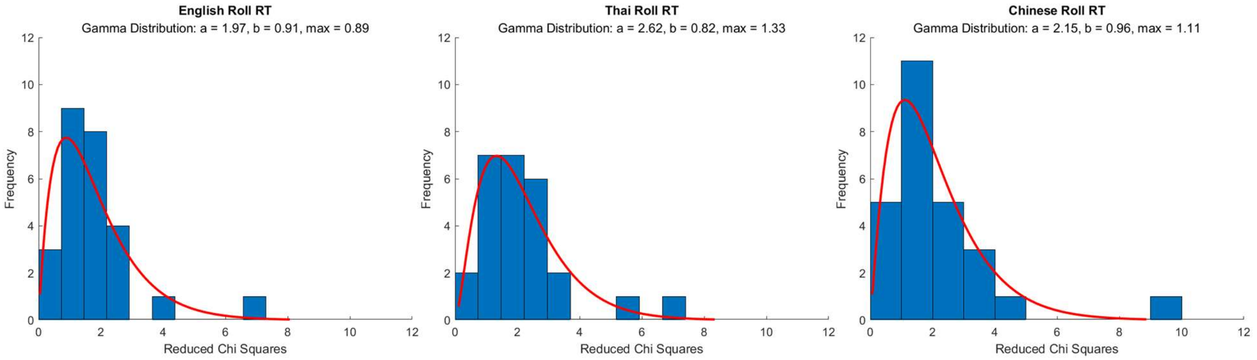


**Figure S4F** Group D distribution of reduced chi squares for individual participant character data. Reduced chi squares for linear fits of both positive and negative angles were analyzed together along with a gamma distribution curve. The data for each character type was divided into 10 bins.

### Supplementary Information

### Figure S5

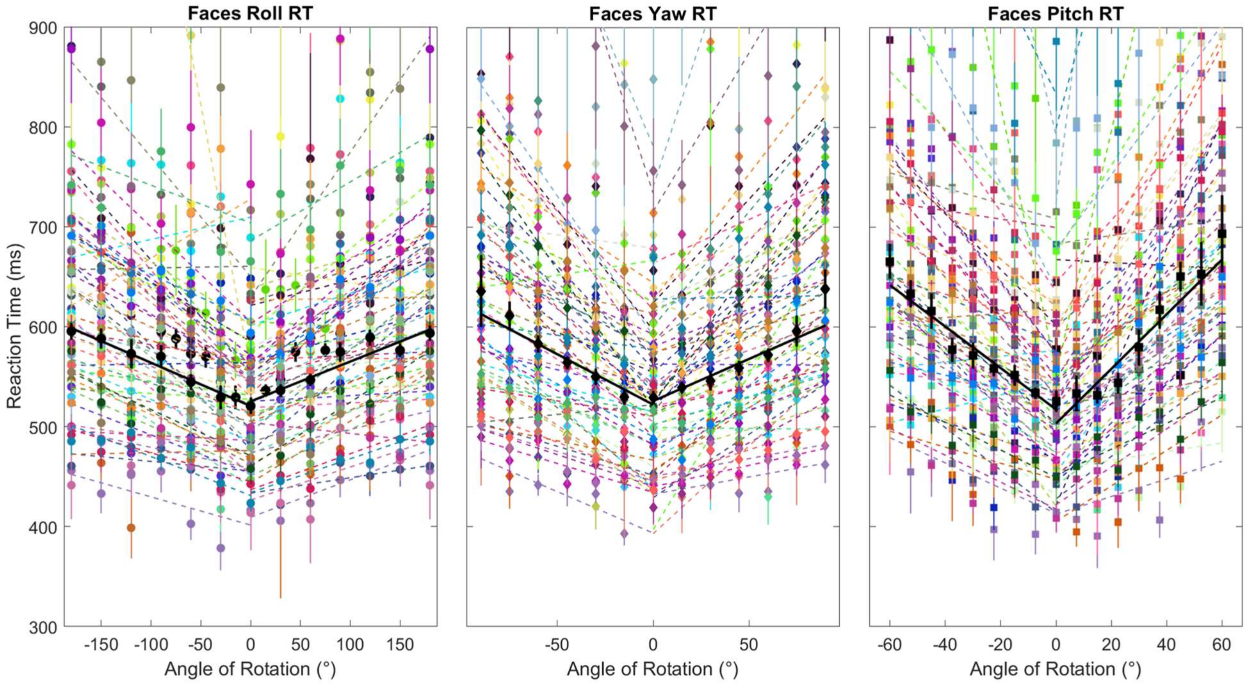


**Figure S5A** Combined pool raw reaction times for faces against roll, yaw, and pitch rotations. Error bars were calculated after a Cousineau normalization was applied.

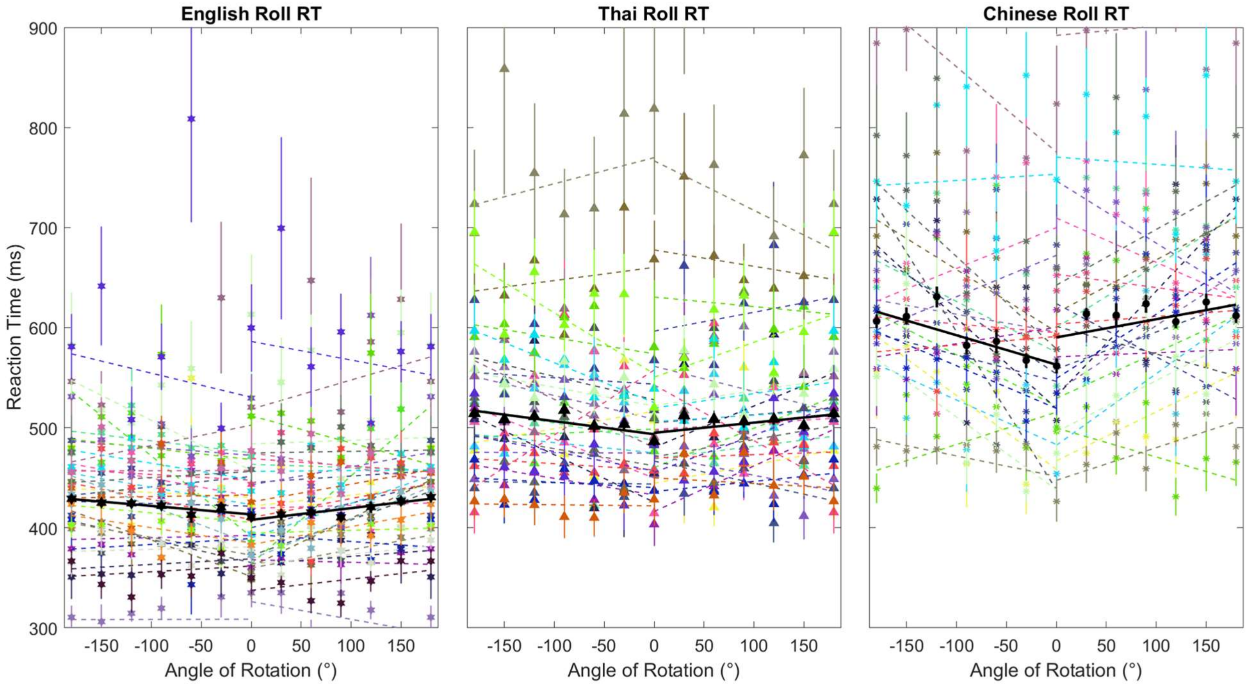


**Figure S5B** Combined pool raw reaction times for English, Thai, and Chinese characters against roll rotation. Error bars were calculated after a Cousineau normalization was applied.

### Supplementary Information

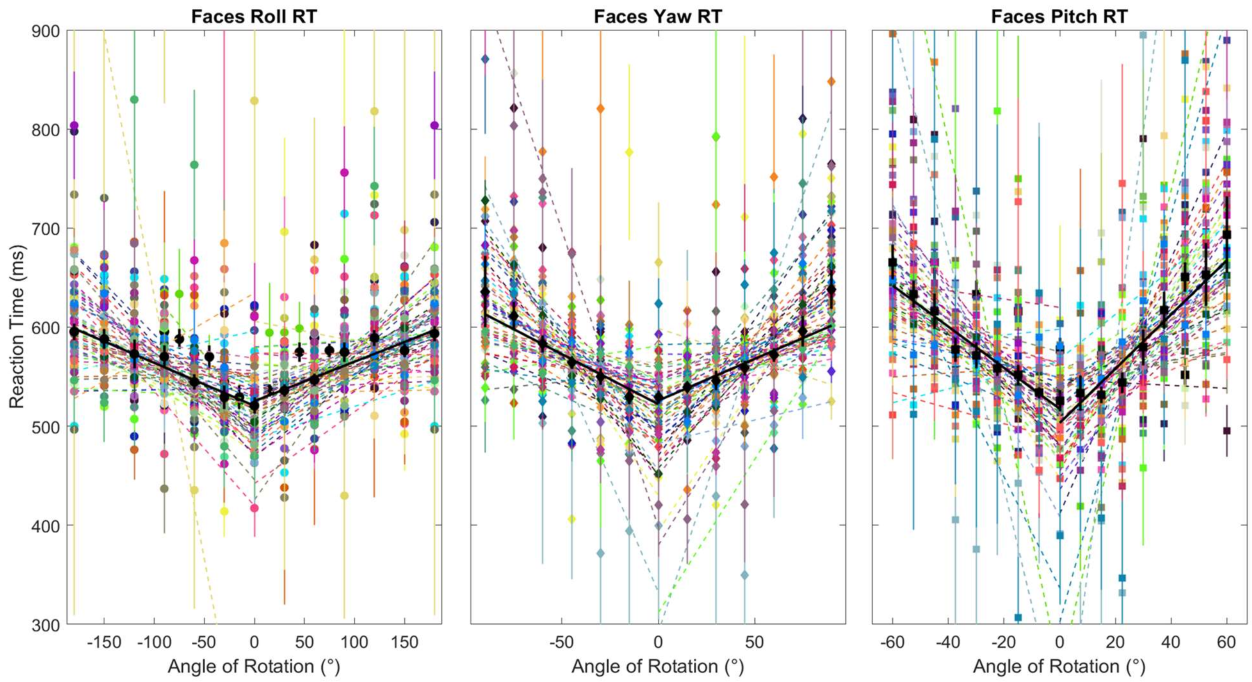


**Figure S5C** Combined data normalized reaction times for faces against roll, yaw, and pitch rotations

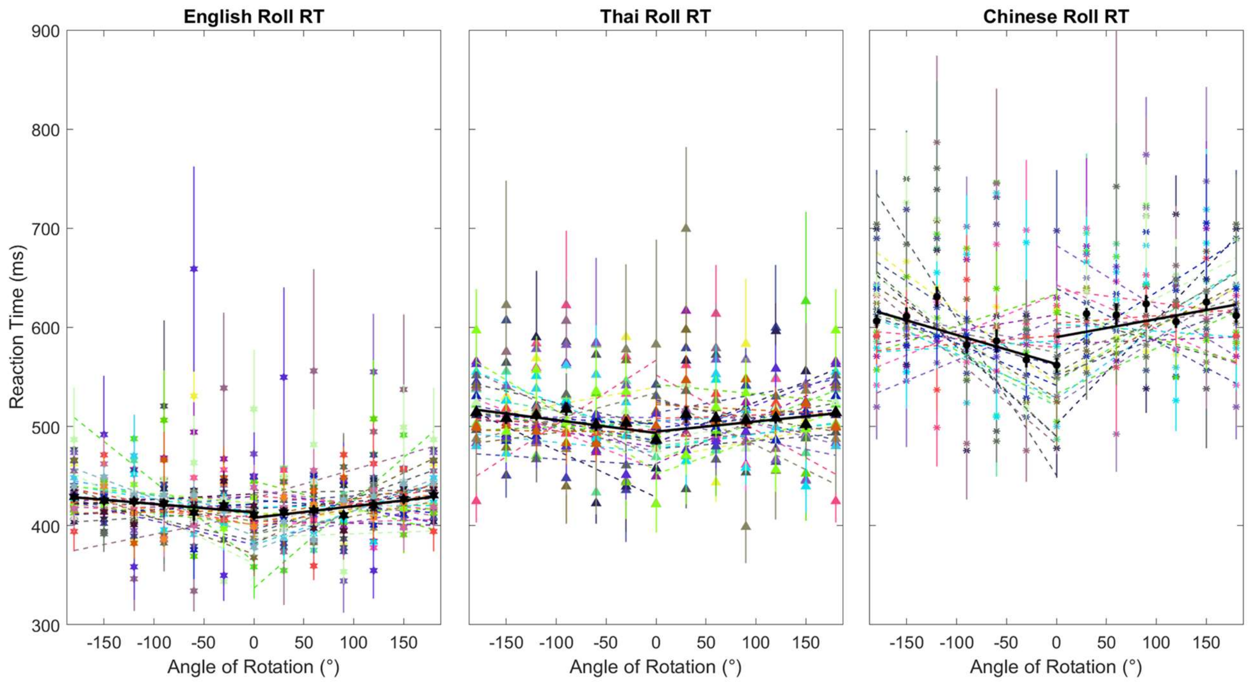


**Figure S5D** Combined data normalized reaction times for English, Thai, and Chinese characters against roll rotation.

### Supplementary Information

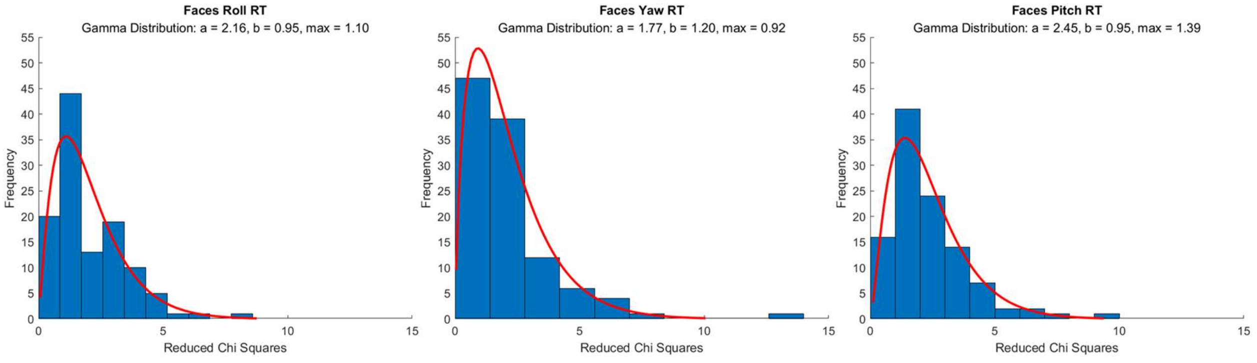


**Figure S5E** Combined pool data distribution of reduced chi squares for individual participant face data. Reduced chi squares for linear fits of both positive and negative angles were analyzed together along with a gamma distribution curve. The data for each character type was divided into 10 bins.

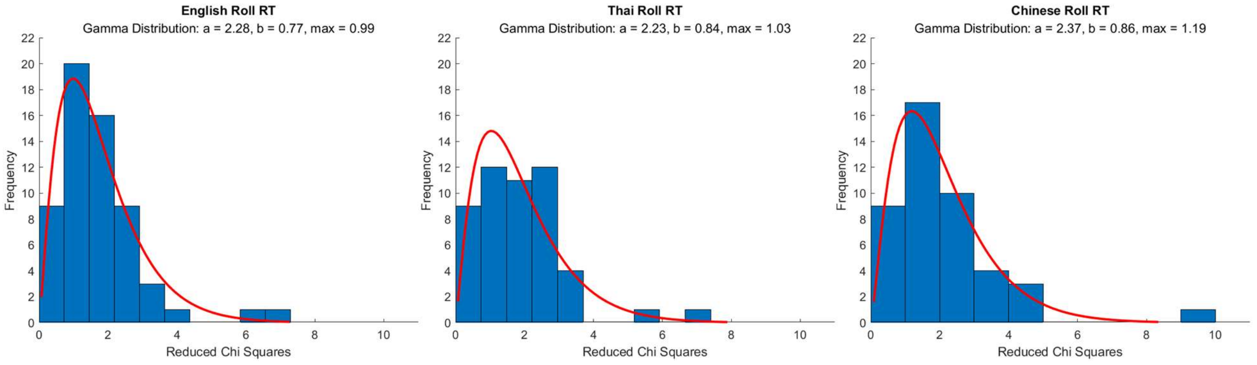


**Figure S5F** Combined pool data distribution of reduced chi squares for individual participant character data. Reduced chi squares for linear fits of both positive and negative angles were analyzed together along with a gamma distribution curve. The data for each character type was divided into 10 bins.
